# Defining the chromatin and transcriptional landscape of stem cell-derived islets

**DOI:** 10.1101/2022.02.26.482126

**Authors:** Punn Augsornworawat, Nathaniel J. Hogrebe, Matthew Ishahak, Erica Marquez, Marlie M. Maestas, Mason D. Schmidt, Daniel A. Veronese-Paniagua, Sarah E. Gale, Julia R. Miller, Leonardo Velazco-Cruz, Jeffrey R. Millman

**Affiliations:** Division of Endocrinology, Metabolism and Lipid Research, Washington University School of Medicine, MSC 8127-057-08, 660 South Euclid Avenue, St. Louis, MO 63110, USA; Department of Biomedical Engineering, Washington University in St. Louis, 1 Brookings Drive, St. Louis, MO 63130, USA

## Abstract

Transplantation of insulin-secreting β-cells differentiated from human pluripotent stem cells holds great potential as a cell therapy for treating insulin-dependent diabetes. While these stem cell-derived islets (SC-islets) are able to reverse diabetes in animal models, they are not fully equivalent to their *in vivo* counterparts. To better define the state of the cell types generated within these SC-islets and provide a resource for identifying deficiencies in lineage specification, we used single-cell multiomic sequencing to simultaneously measure the chromatin accessibility and transcriptional profiles of SC-islets at multiple time points as well as primary human islets. The integrated analysis of both the transcriptional and chromatin landscape for each cell provided greater resolution for defining cell identity, allowing us to derive novel gene lists for identifying each islet cell type. Furthermore, this multiomic analysis revealed that the difference between SC-β cells and enterochromaffin-like cells, which are a major off-target from *in vitro* differentiation, is a gradient of progressive cell states rather than a stark difference in identity. The chromatin landscape of primary human islets was much more restricted, suggesting that stem cell-derived cells are not fully locked into their cell fate. While long term culture of SC-islets both *in vitro* and *in vivo* does close overall chromatin state, only *in vivo* transplantation directs cells toward their correct identities. Collectively, our multiomic analysis demonstrates that both the chromatin and transcriptional landscapes play significant roles in islet cell identity, and these data can be used as a resource to identify specific deficiencies in the chromatin and transcriptional state of SC-islet cell types.

## Introduction

The development of methods to differentiate human pluripotent stem cells (hPSCs) into islet-like clusters has the potential to generate an unlimited number of insulin-producing stem cell-derived β (SC-β) cells for the treatment of insulin-dependent diabetes ^1-3^. This process utilizes temporal combinations of small molecules and growth factors ^4-6^, microenvironmental cues ^7^, and other sorting or aggregation ^4, 8-11^ approaches to drive cells through several intermediate progenitor cell types. The resulting SC-β cells possess many features of primary human β-cells, including the expression of β-cell specific markers, glucose responsive insulin secretion, and the ability to reverse severe diabetes in animal models ^5, 6, 12-15^. As a result, these cells have garnered great interest and potential to provide a functional cure for human patients with type 1 diabetes. However, these stem cell-derived islets (SC-islets) are a heterogeneous tissue that also contains stem cell-derived α (SC-α) and δ (SC-δ) cells, as well as an endocrine cell type of intestinal lineage denoted here as stem cell-derived enterochromaffin-like (SC-EC) cells ^16^. The presence of this off-target population suggests that there are inefficiencies in lineage specification during these directed differentiation protocols ^9^. Correcting this aberrant signaling could enhance specification to a β-cell identity and further improve the function of SC-β cells.

The specific pattern of gene expression and chromatin state within a cell directs differentiation and maintains its final identity ^17-20^. Thus, characterizing both the transcriptional and chromatin landscape of cells during this differentiation process can provide insight into the degree that a particular cell type resembles its *in vivo* analogue. Single-cell RNA sequencing has been used to characterize the transcriptional environment of SC-islets, demonstrating that SC-β cells express many, but not all, of the important genes found in human primary β-cells ^9, 13, 14, 21, 22^. Studies have also begun to investigate the chromatin state within pancreatic cell differentiations using bulk approaches ^23-25^, and recent work has demonstrated the advantages of studying primary human islets using single-cell Assay for Transposase-Accessible Chromatin (ATAC) sequencing ^26-28^. ATAC sequencing can describe whether particular chromatin regions are in an open, accessible state that is ready to be transcribed or interacted with, providing valuable information about cell identity that is missing when only transcriptional data is investigated. This data can define, for instance, the chromatin accessibility of promoters for the specific genes themselves that are capable of being transcribed or the DNA sequence motifs where specific transcription factors bind to help activate the transcription of different genes. Importantly, modulating the chromatin states of particular genes and motifs that are mis-expressed in *in vitro*-derived cell types are likely to drive cells closer to the identity of their *in vivo* counterparts. Despite the potential power of using a single-cell multiomic approach, a comprehensive indexing of the combined chromatin accessibility and transcriptional signatures of each *in vitro*-differentiated SC-islet cell type is lacking.

Here, we provide a multiomic analysis of the transcriptional and chromatin landscapes of SC-islets at the single-cell level. The simultaneous characterization of both the transcriptional and chromatin states for each cell allowed us to better define each cell type found within SC-islets, resulting in novel lists of important genes and motifs for each cell type. Interestingly, SC-EC and SC-β cells formed a gradient of cell identities rather than distinct cell types, with sub-populations of each cell type exhibiting characteristics of the other. Primary human islets had a much more restricted and defined chromatin state, while SC-islet cell types had open chromatin regions that were associated with other lineages. Transplantation into mice for 6 months closed many of these accessible chromatin regions and improved the expression of signature genes associated with each lineage, while extended *in vitro* culture did not have the same effect on identity. Furthermore, we identified and modulated chromatin regulators that were important for SC-β cell identity, highlighting the significance of the chromatin landscape. Collectively, these data provide a resource that improves the characterization of cell identities found within SC-islets, facilitating the discovery of mis-expressed genes and motifs that can be targeted to further improve SC-β cell differentiation strategies.

### Multiomic characterization of SC-islets enables greater resolution of cell identity

To better understand the cell types produced with directed differentiation strategies toward pancreatic islets, we sought to use an integrated analysis of both the transcriptional and chromatin landscapes for each cell to define cell identity and ascertain deficiencies in lineage specification more accurately. We differentiated human pluripotent stem cells (hPSCs) through our 6-stage protocol using specific temporal combinations of small molecules and growth factors to produce islet-like clusters ^7, 29^ (Extended Data Fig. 1a and b, Supplementary Table 1). These SC-islets were processed and sequenced using single-cell multiomics, obtaining both gene expression (mRNA) and chromatin accessibility (ATAC) information for each cell. (Fig. 1a, Supplementary Table 2). Analysis of gene expression and chromatin accessibility both individually and in combination enabled us to identify specific islet cell types, including SC-β, SC-α, and SC-δ cells as well as the off-target populations SC-EC cells, exocrine, and mesenchymal-like cells (Fig. 1b and c, Extended Data Fig. 1c and d). While both gene expression and chromatin accessibility information on their own were sufficient to separate most cell types in SC-islets, we were able to identify two subpopulations of SC-EC cells (denoted as SC-EC1 and SC-EC2) using the integrated analysis of both mRNA and ATAC data (Fig. 1b). These EC populations clustered adjacent to SC-β cells, suggesting their resemblance. In contrast, SC-α and SC-δ populations formed more distinct, separated clusters (Fig. 1b).

**Fig 1.**
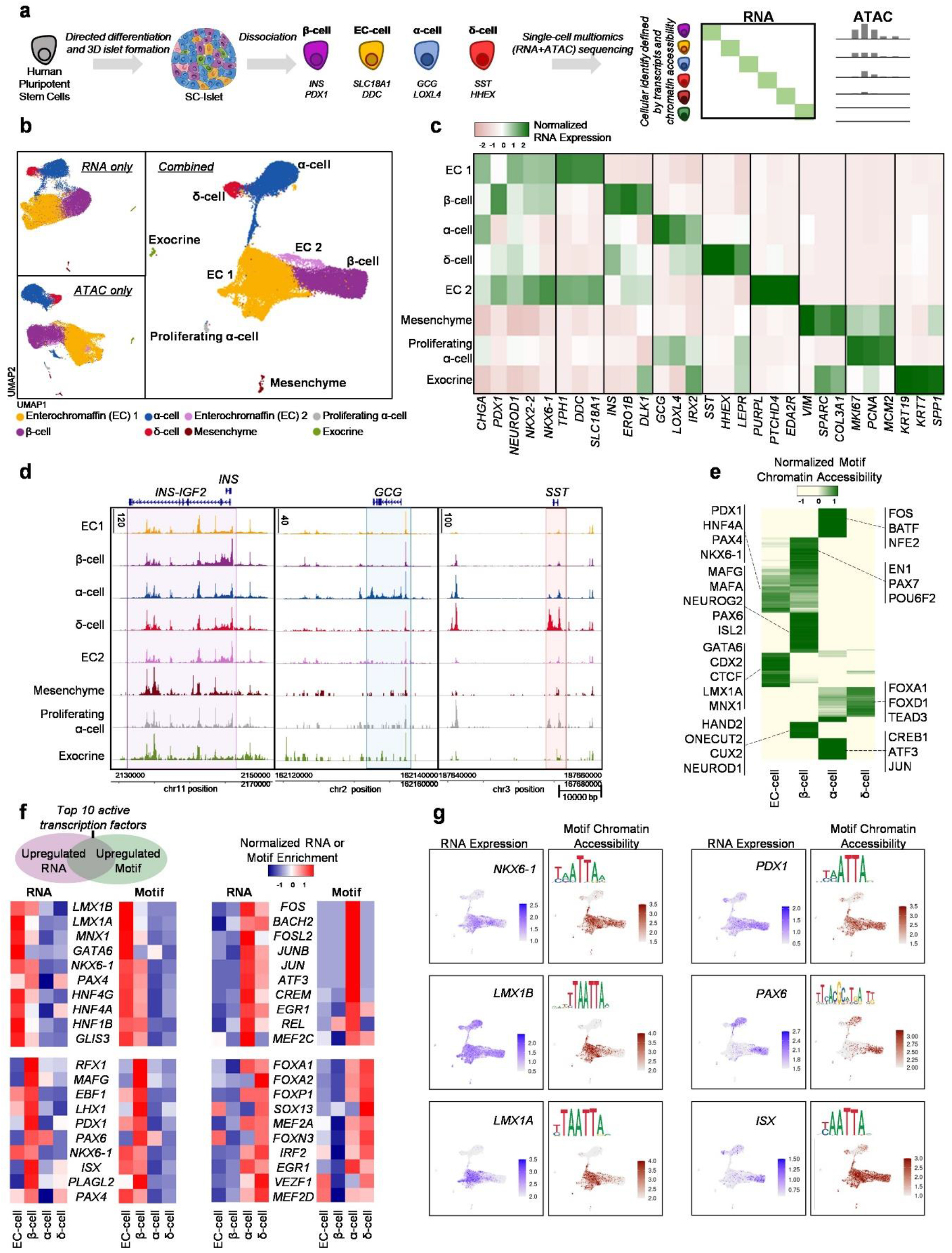
Multiomic profiling of stem-cell derived islets shows unique chromatin accessibility signatures in endocrine cell types. a, Schematic of SC-islet differentiation and multiomic sequencing. b, UMAPs showing identified cell types in SC-islets 2 weeks into stage 6 using both and either chromatin accessibility (ATAC) or gene (mRNA) information. c, Heatmap showing gene expression of markers associated with each cell type. d, ATAC plots showing chromatin accessibility of SC-islet cell types around the INS, GCG, and SST genomic regions. e, Heatmap showing the top 200 variable DNA-binding motif accessibility within endocrine cell populations and highlighting markers for each cell type. f, Heatmaps highlighting gene expression and ATAC motif accessibility of top 10 active transcription factors co-enriched with both features in SC-β, SC-α, SC-δ, and SC-EC cells. g, UMAP showing gene expression and motif accessibility of selected transcription factors associated with SC-β cells or SC-EC cells. SC, Stem cell derived; EC, enterochromaffin

Each of the identified populations had high expression of genes traditionally associated with their cell type (Fig. 1c). Furthermore, the SC-β, SC-α, and SC-δ populations had strong chromatin peaks represented by their respective *INS, GCG*, and *SST* gene regions as expected (Fig. 1d). Unexpectedly, however, the *INS* gene region had open accessibility across all detected cell types, including non-endocrine cell types to some extent. In contrast, *GCG* and *SST* gene regions had relatively few chromatin peaks outside of SC-α cells and SC-δ cells, respectively. We also found distinct motif groups enriched in each specific cell type (Fig. 1e, Extended Data Fig. 1e, Supplementary Table 4). Within the endocrine population, SC-β cells had enrichment of motifs that correspond to known β-cell associated transcription factors ^9, 22, 30, 31^. Interestingly, this list includes the DNA-binding motif for MAFA, which is a transcription factor associated with primary β-cells but is expressed at a very low level in SC-β cells ^16, 32^. Motifs that were specifically enriched in SC-EC cells include GATA6, CDX2, CTCF, and LMX1A, which are often associated with intestinal lineages ^33, 34^. Interestingly, SC-β cells and SC-EC cells possessed shared enrichment of certain motifs, such as PDX1, PAX4, and NKX6-1, which have been reported to be important in human primary β-cells ^27^ (Fig. 1e).

In an effort to better characterize these cell populations within SC-islets, we cross-referenced transcription factor DNA-binding motif chromatin accessibility and gene expression information to identify active transcription factors that have both high expression and that can access their target binding motifs to promote transcription of other genes (Supplementary Table 4). Here, we highlight the top 10 identified active transcription factors enriched in specific endocrine populations compared to the average of the other endocrine populations (Fig. 1f). Highly active transcription factors in SC-β cells include of RFX1, PDX1 and PAX6, which have been previously reported to be important for β-cell identity ^27, 35^. Other highly active transcription factors, such as MAFG, EBF1, ISX, and PLAGL2, have not been highlighted in previous studies, demonstrating the utility of using both mRNA and chromatin accessibility data for identifying cell types. In contrast, SC-EC cells have co-enrichment of LMX1B, LMX1A, MNX1, and GATA6. Notably, the SC-β and SC-EC populations both have high activities of NKX6-1 and PAX4, suggesting that they share common features essential for their identities. This similarity is further demonstrated with motif chromatin accessibility plots, where NKX6-1 and PDX1 were enriched in both SC-β and SC-EC populations (Fig. 1e and g). In contrast, binding motifs for PAX6 were enriched in SC-β cells compared to SC-EC cells while LMX1A was enriched in SC-EC cells compared to SC-β cells. Unexpectedly, we found a mismatch in RNA expression and motif accessibility across cell types for certain transcription factors (Fig. 1f and g). In one instance, a high amount of *LMX1B* transcript was detected in all endocrine cell types, but motif accessibility was predominately open only in SC-EC cells, indicating that this transcription factor is likely helping activate transcription of its target genes only in SC-EC cells. In contrast, *ISX* transcripts were detected predominately in SC-β and SC-δ cells, but the associated DNA-binding motif was enriched in SC-β and SC-EC cells. Thus, while the motif is open in SC-EC cells, only SC-β cells are actively transcribing the transcription factor needed to utilize this open motif. Collectively, this multiomic analysis has generated novel lists of genes to better classify the cell types found within SC-islets, providing better resolution of cell identity and insights into the relative importance of different transcription factor activity within each cell type.

### SC-EC and SC-β cells form a gradient of progressive cell states rather than well-defined cell populations

SC-EC cells are an off-target cell population that arises during *in vitro* SC-islet differentiation protocols but that does not positively contribute to tissue function ^9, 36, 37^ (Fig. 1b). Although both enterochromaffin and β-cells arise from a definitive endoderm precursor during *in vivo* development, intestinal and pancreatic origins respectively, they appear to share a common progenitor lineage during *in vitro* differentiation of SC-islets ^9^. However, the field does not fully understand the nature of these cell populations that emerge from differentiation. Interestingly, the combined mRNA/ATAC clustering analysis allowed us to resolve two distinct SC-EC populations that were adjacent to the SC-β cell population (Fig. 1a), suggesting that there may be multiple or a gradient of cell states between SC-EC and SC-β cells. Given the apparent similarity of SC-β and the two SC-EC cell populations in our multiomic clustering (Fig. 1b), we performed a trajectory analysis to detect differences in both gene expression and chromatin accessibility between SC-β and SC-EC cells (Fig. 2a, Extended Data Fig. 2a, Supplementary Table 5). Enterochromaffin cell identity genes, such as *SLC18A1, TPH1*, and *FEV*, were most highly expressed in the SC-EC side of the trajectory map as expected. Similarly, β-cell identity genes, including *INS, PAX6*, and *ISL1*, were most highly expressed toward the SC-β cell end (Fig. 2b and c). Furthermore, gene sets relating to SC-EC or SC-β cells were upregulated in their respective sides of the trajectory map (Extended Data Fig. 2b). Differential motif chromatin accessibility analysis identified LMX1A, GATA6, and CTCF motifs to be enriched in the SC-EC cell end, while NEUROD1, HAND2, MAFA, PAX6, ONECUT2, and RFX2 motifs were enriched for in the SC-β cell side (Fig. 2b and c).

**Fig 2.**
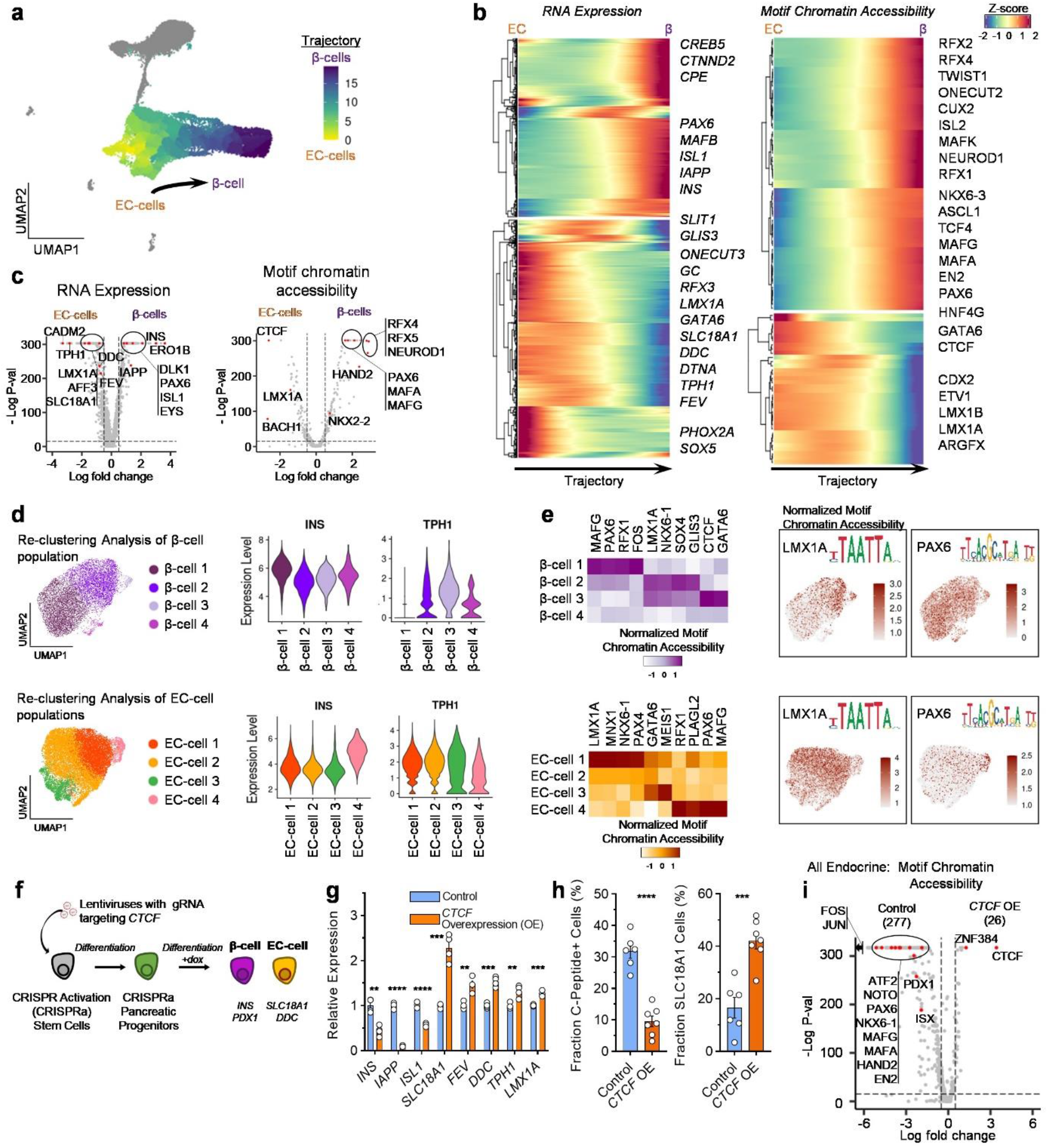
SC-EC and SC-β cells have unique and common transcriptional and chromatin accessibility signatures. a, UMAP showing the trajectory of cells from the SC-β, SC-EC1 and SC-EC2 population. b, Trajectory heatmap showing dynamic changes of gene expression and motif accessibility enriched in β, and EC groups. Selected genes and motifs that are relevant to SC-β, and SC-EC identity and development are highlighted. c, Volcano plots showing differential gene expression analysis (left) and differential motif accessibility analysis (right), highlighting relevant genes associated with SC-β and SC-EC cell populations. d, UMAP showing sub-populations by reclustering the SC-β population or SC-EC cell populations. Violin plots show gene marker expressions of *INS* and *TPH1*, highlighting the presence of off-target genes in sub-populations. e, Heatmap showing DNA-binding motif accessibility associated with SC-β and SC-EC in subpopulations. Selected transcription factors are plotted to show distribution of cells with target or off-target motif accessibility. f, Schematic of CRISPRa experiment for overexpression of *CTCF* in differentiating pancreatic progenitor cells. g, qPCR analysis of differentiated SC-islets with *CTCF* over expression during endocrine induction, plotting mean ± s.e.m. (n = 4), showing expression differences of genes associated with β-cells (*INS*, P = 0.0010; *IAPP*, P = 1.5 × 10^−7^; *ISL1*, P = 3.0 × 10^−5^) and EC cells (*SLC18A1*, P = 1.6 × 10^−4^; *FEV*, P = 0.0019; *DDC*, P = 1.2 × 10^−4^; *TPH1*, P = 0.0047; *LMX1A*, P = 4.6 × 10^−4^). Statistical significance was assessed by unpaired two-sided t-test. h, Immunocytochemistry quantification of cells expressing C-peptide protein (P = 5.3 × 10^−6^) and SLC18A1 protein (P = 3.1 × 10^−4^) with or without *CTCF* over expression, plotting mean ± s.e.m. (control; n = 6, doxycycline; n = 7). Statistical significance was assessed by unpaired two-sided t-test. i, Volcano plots showing differential motif chromatin accessibility analysis of SC-endocrine population comparing control and *CTCF* over expression. SC, Stem cell derived; EC, enterochromaffin cells; CRISPRa, CRISPR activation.

While many of these identified genes and motifs were enriched on the side normally associated with that particular cell type, their expression was often a gradient that spilled over toward the other end of the trajectory map (Fig. 1b). To further probe the transcriptional and chromatin landscape of cells along this trajectory map, we performed subclustering analyses separately on the SC-β cell population and the SC-EC cell populations (Fig. 2d). This subclustering identified 4 separate sub-populations of both SC-β and SC-EC cells (Fig. 2d). Differential expression analyses revealed that each of the SC-β cell sub-populations had distinguishing gene expression and motif accessibility features (Fig. 2d and e, Extended Data Fig. 2c and d, Supplementary Table 6), including SC-β cells with *TPH1* expression and more open chromatin accessibility of the SC-EC cell-associated DNA-binding motifs for LMX1A and CTCF. The high *INS*-expressing SC-β cell population displayed greater motif accessibility to β-cell-associated transcription factors, such as PAX6, and less motif accessibility to EC-cell transcription factors, such as LMX1A. Similarly, differential expression analyses demonstrated that the four SC-EC cell sub-populations also had distinguishing gene expression and motif accessibility features, including SC-EC cells with elevated *INS* expression and more open chromatin accessibility of the SC-β cell-associated DNA-binding motifs for PAX6 and MAFG. Collectively, these data suggest that the SC-EC and SC-β cell populations produced from *in vitro* differentiation form a continuum of cell states rather than exhibiting clear exclusivity of gene expression and chromatin accessibility. The identification of cells with both SC-β and SC-EC cell features here is in contrast with prior studies that only used transcriptional profiling of SC-islets to conclude that the SC-β and SC-EC cells populations were distinct ^9, 12^, highlighting how this multiomic approach facilitates greater resolution of cell identity.

We were interested in understanding how chromatin regulators could influence cell fate identities between islet and intestinal endocrine cells during *in vitro* differentiation. Our integrated multiomic analysis we identified the chromatin remodeler CCCTC-binding factor (CTCF) as the transcription factor binding motif having the greatest increase in accessibility in SC-EC cells compared to SC-β cells (Fig. 2c). Because CTCF is a chromatin remodeler that is involved in other developmental and differentiation processes ^38^, we sought to further examine its impact on pancreatic differentiation. We utilized a doxycycline inducible VP64-p65-Rta CRISPR activation (CRISPRa) stem cell line ^39^ to increase transcription of *CTCF* during differentiation (Fig. 2f). Unfortunately, we were unsuccessful in also knocking down its expression despite trying several approaches, which we would hypothesize could improve differentiation to SC-β cells. The introduction of *CTCF* guide RNA (gRNA) into the CRISPRa stem cell line induced the upregulation of *CTCF* expression upon activation of *dCas9* with doxycycline (Extended Data Fig. 3a). Upregulation of *CTCF* during stage 5 endocrine cell induction ^7, 29^ resulted in drastic reductions in the expression of β-cell identity genes, such as *INS* and *ISL1*, and significant upregulation of intestinal lineage associated genes, such as *SLC18A1* and *FEV* (Fig. 2g, Extended Data Fig. 3a). At the end of differentiation, *CTCF*-overexpressed cells displayed upregulated EC-cell genes including *SLC18A1, FEV, DDC, TPH1*, and *LMX1A* (Extended Data Fig. 3b). Immunocytochemistry staining revealed a greatly diminished SC-β cell population but increased SC-EC cell generation (Fig. 2h, Extended Data Fig. 3c). This change in lineage was further demonstrated with reduced total insulin content and impaired glucose-stimulated insulin secretion in *CTCF*-overexpressed cells (Extended Data Fig. 3d and e). The effects of *CTCF* overexpression were temporally dependent and had the greatest effect during the endocrine induction stage of the protocol, suggesting *CTCF* to be important for SC-EC versus SC-β cell fate selection (Extended Data Fig. 3f). Furthermore, single-cell multiomic sequencing demonstrated that CTCF overexpression caused endocrine cells to have increased CTCF motif chromatin accessibility and SC-EC cell signatures while decreased accessibility of several β-cell-associated transcription factor binding motifs, such as PAX6, MAFA, PDX1, and ISX (Fig. 2i, Extended Data Fig. 3g to l). Taken together, these findings show that *CTCF* modulation drastically disrupted acquisition of a SC-β cell identity from pancreatic progenitors and instead redirected them to an intestinal enterochromaffin-like cell fate, highlighting the utility of this multiomics approach for identifying important regulators of SC-β cell fate determination.

**Fig 3.**
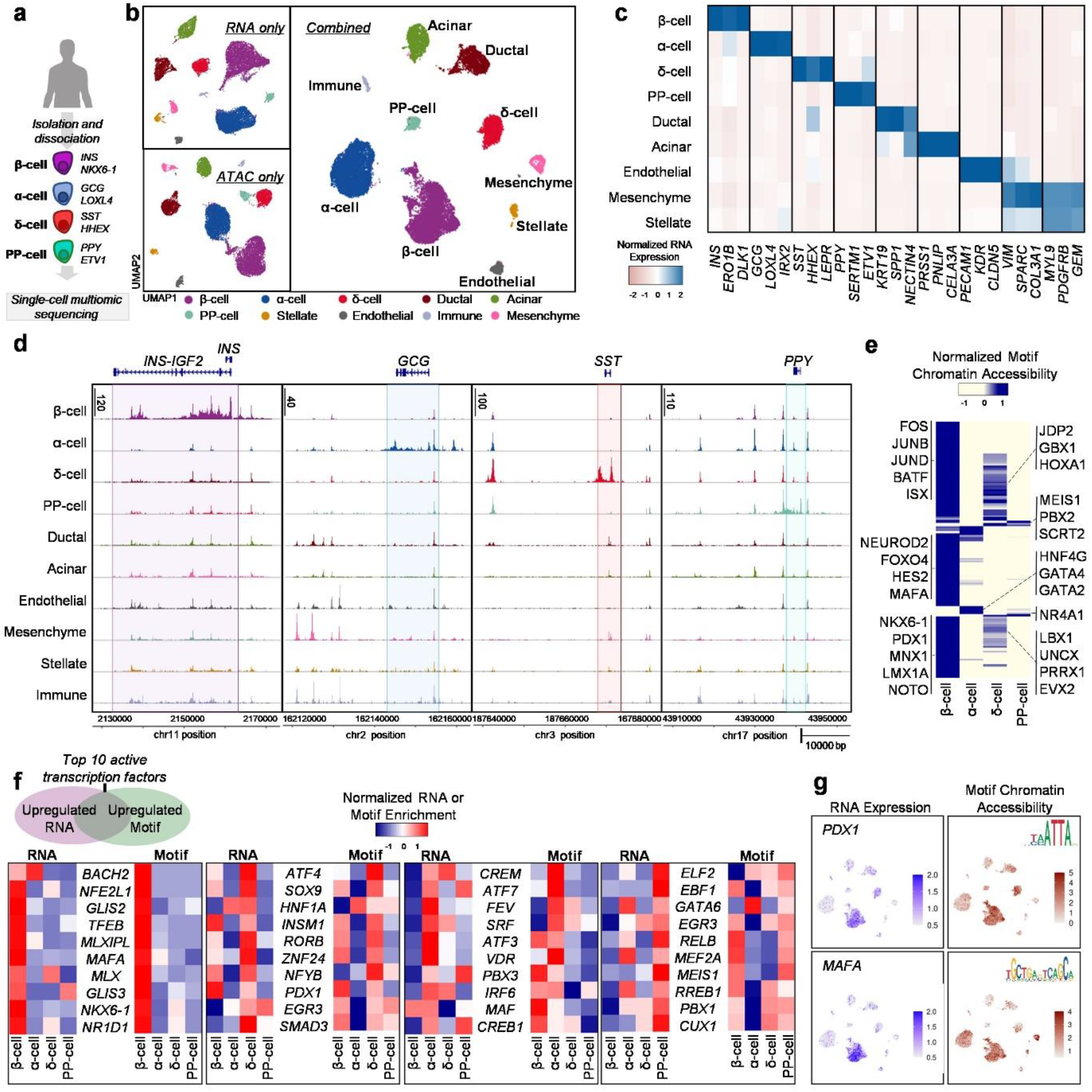
Multiomic profiling of human adult primary islets shows unique chromatin accessibility signatures in endocrine cell types. a, Schematic of human adult primary islets for mutiomic sequencing. b, UMAPs showing identified cell types in primary human islets using both and either chromatin accessibility (ATAC) or gene (mRNA) information. c, Heatmap showing gene expression of markers associated with each cell type. d, ATAC plots showing chromatin accessibility primary islet cell types around the *INS, GCG, SST*, and *PPY* genomic regions. e, Heatmap showing the top 200 variable DNA-binding motif accessibility within endocrine cell populations and highlighting motif markers for each cell type. f, Heatmaps highlighting gene expression and ATAC motif accessibility of top 10 active transcription factors co-enriched with both features in primary β, α, δ, and PP-cells. g, UMAP showing gene expression and motif accessibility of selected transcription factors associated with primary β-cells. PP, pancreatic polypeptide.

### Multiomic characterization of primary islets reveals well-defined cell types

In order to compare the transcriptional and chromatin signatures identified in our SC-islets to their *in vivo* counterparts, we sequenced and characterized primary human islets using the same multiomics approach (Fig. 3a, Extended Data Fig. 4a and b, Supplementary Table 2). Using mRNA and chromatin information either in isolation or in a combined analysis, we identified ten distinct populations that included both pancreatic endocrine and exocrine cell types as well as other minor cell populations (Fig. 3b and c, Extended Data Fig. 4c and d, Supplementary Table 3). Unlike SC-islets (Fig. 1b and 1c, Extended Data Fig. 1c). Clusters representing all cell types appear very distinct and exhibit robust expression of their respective gene markers by. In addition, the chromatin accessibility profiles demonstrate that primary islet endocrine cells contain distinct peak signals around the *INS, GCG, SST*, and *PPY* genomic regions of β-cells, α-cells, δ-cells, and PP-cells, respectively (Fig. 3d). The closed chromatin accessibility profile of *INS* across non-β-cell types in particular is in stark contrast to that observed in SC-islets (Fig. 1d), where chromatin accessibility of the *INS* gene is relatively open across all cell types. We also performed chromatin motif accessibility analyses in primary islets and identified distinguishing features across these endocrine types (Fig. 3e, Extended Data Fig. 4e, Supplementary Table 4). Notably, primary β-cells have accessible binding sites for PDX1, NKX6-1, MAFA and ISX, similar to SC-β cells. Surprisingly, however, motifs identified as enriched in SC-EC cells, such as LMX1A and MNX1, were enriched in β-cells when compared to other endocrine populations. Furthermore, re-clustering analysis of the primary β-cells identified 3 sub-populations that displayed unique gene expression and accessible motif signatures (Extended Data Fig. 4f - j, Supplementary Table 6), consistent with previous studies utilizing either single-cell RNA or ATAC sequencing ^27, 40-42^.

**Fig 4.**
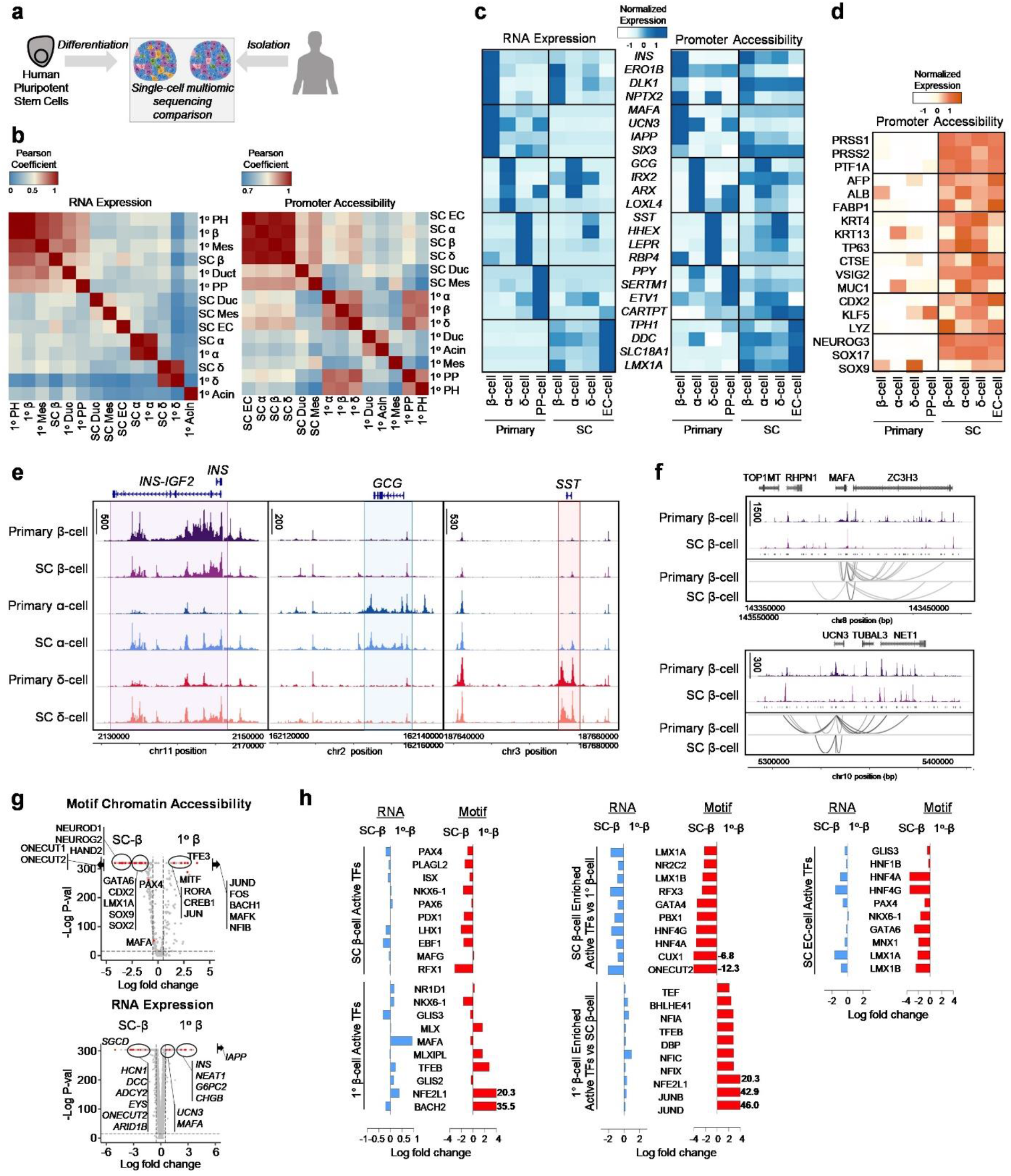
Comparative analysis of SC-islets and primary human islets shows differences in chromatin accessibility signatures associated with islet identity. a, Schematic showing comparisons of SC-islet and primary human islets. b, Pearson correlation analysis comparing cell types in SC-islets and primary islets using gene expression and ATAC promoter accessibility. c, Heatmap highlighting and comparing identity (β, adult β, α, δ, PP and EC) associated gene expression and ATAC promoter accessibility of endocrine cells in SC-islets and primary islets. d, Heatmap highlighting and comparing off-target identity (Exocrine, hepatic, esophagus, stomach, intestinal, pancreatic progenitor) associated gene expression and ATAC promoter accessibility in SC-islet cells and primary islet cells. e, ATAC plots comparing chromatin accessibility around the *INS, GCG*, and *SST* genomic regions in β, α, and δ-cells from SC-islets and primary islets. f, ATAC peaks from SC-β cells and 1° β-cells showing chromatin accessibility around β-cell identity marker, *MAFA* (top) and *UCN3* (bottom). Peaks were linked and analyzed to depict differences in the number of cis-regulatory elements. c, Volcano plots showing differential motif accessibility analysis (top) and differential gene expression analysis (bottom) comparing SC-β cells and primary β-cells. h, Bar graphs showing fold change differences between SC-β cells and primary β-cells, showing gene expression and motif accessibility of identified transcription factors associated with SC-β cells, primary β-cells, and SC-EC cells. SC, stem cell derived; EC, enterochromaffin cells.

Similar to our multiomic analysis of SC-islets, we also examined transcription factor activity in primary islet endocrine cells as assessed by both relative increases in mRNA transcripts and chromatin accessibility of the corresponding DNA-binding motif (Fig. 3f and g). For β-cells, NKX6-1 is the only transcription factor that is shared in the top 10 active factor list with SC-β cells (Fig. 1f vs Fig 3f). Other notable active transcription factors include TFEB, MLX, NR1D1 and the primary β-cell gene *MAFA*. δ-cells share multiple accessible motifs with β-cells, including ATF4, SOX9, INSM1, and PDX1. NFYB, EGR3, and SMAD3, however, are exclusive to δ-cells by both motif enrichment and mRNA. Notable α-cell active factors include CREM, ATF7, ATF3, and VDR, while PP-cells were enriched for ELF2, EBF1, MEIS1, and CUX1. While some of the active transcription factors are shared with the list from SC-islets, this analysis illustrates that primary islet cells have a unique transcriptional and chromatin landscape when compared to their stem cell-derived counterparts, highlighting specific deficiencies in lineage specification during directed differentiation protocols.

### Primary human islets have a more restricted chromatin landscape than SC-islets

Previous studies have demonstrated that β-cells derived from *in vitro* differentiation of hPSCs are functionally and transcriptionally different than their *in vivo* counterparts ^5, 6, 9, 13, 14, 16^. To obtain greater resolution of the differences between stem cell-derived endocrine cells and those found *in vivo*, we used single-cell multiomics to directly compare both the transcriptional and chromatin landscapes of SC-islets and primary human islets (Fig. 4a). Both SC-islets and primary islets contained β, α, and δ-cells (Extended Data Fig. 5a and b). However, only SC-islets contained EC-cells, while only primary islets contained PP-cells, consistent with prior analysis ^9^. Pearson correlation analysis of gene expression demonstrated that SC-islet cell types were generally most similar to their primary cell counterparts (Fig. 4b). Surprisingly, this trend was not observed with promoter chromatin accessibility. Greater similarity was observed based on cell origin (*in vitro*-derived vs. primary) rather than by cell type. We explored this dissimilarity in promoter accessibility further by examining commonly used endocrine identity genes (Fig. 4c). In general, promoter accessibility of SC-islet cells was broadly open across cell types even though mRNA expression patterns were consistent with prior RNA-only analysis ^5, 9, 13^. SC-β cells expressing high levels of many identity genes but yet maintaining low expression of *MAFA, UCN3, IAPP*, and *SIX2* (Fig. 4c). Additionally, SC-islet cells have open chromatin at gene regions associated with non-endocrine identities including exocrine, hepatic, esophagus, stomach, intestinal, and progenitor cells (Fig. 4d). Notably, primary β-cells had greater chromatin accessibility of the *INS* gene compared to SC-β cells (Fig. 4e). SC-α and δ-cells also shared chromatin accessibility peaks around the *INS* gene, which was not observed in primary α and δ-cells. The *GCG* and *SST* gene regions were open in α and δ-cells from both origins, respectively. Unlike with *INS*, however, these gene regions were relatively inaccessible in the other endocrine cell types, regardless of origin. Further analysis of the chromatin accessibility of *MAFA* and *UCN3* (Fig. 4f), which are known to be transcriptionally upregulated in β-cells ^9, 13, 43-45^, demonstrated that predicted cis-regulatory elements present around the *MAFA* and *UCN3* gene regions differed considerably between SC-β and primary β-cells.

**Fig 5.**
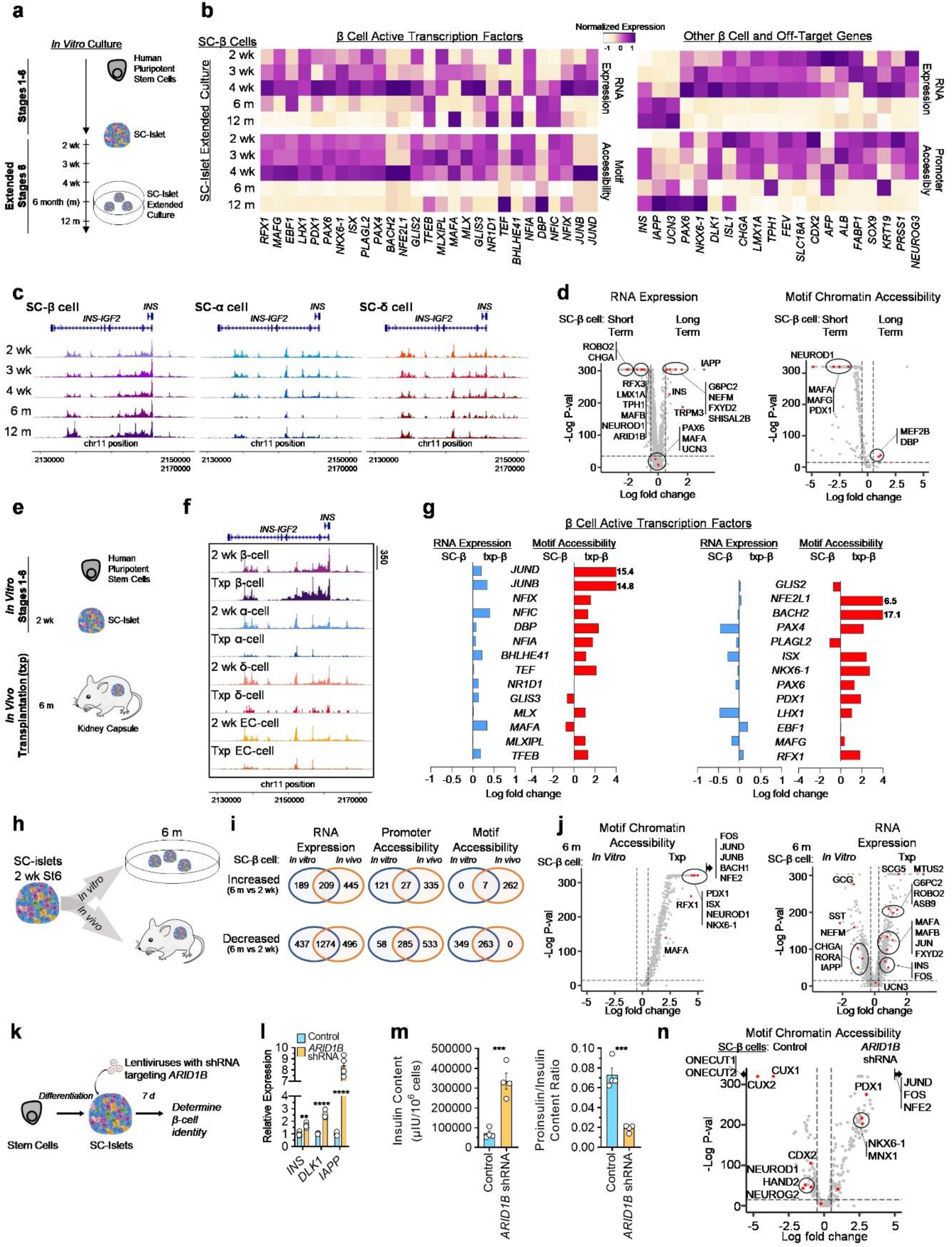
Interrogating SC-islet identity with time *in vitro* and after transplantation. a, Schematic of SC-islet *in vitro* culture with extended time course. b, Heatmap showing gene expression, motif chromatin accessibility, or ATAC promoter accessibility of gene markers and transcription factors of SC-β cells cultured *in vitro* with extended time (2, 3, 4 weeks and 6, 12 months). c, Graphs comparing time course of chromatin accessibility around the *INS* genomic region in SC-β, α, and δ cells. Peak signals around the *INS* gene in SC-β cell increases over time, but decreases in SC-α, and δ cells. d, Volcano plots showing differential gene expression analysis (left) and differential motif accessibility analysis (right) comparing SC-β cells cultured short term (week 2, 3 and 4) and SC-β cells cultured long term (month 6 and 12). e, Schematic of differentiated SC-islets maintained long-term by *in vivo* transplantation. f, ATAC chromatin accessibility around the *INS* genomic region showing increase of peak signals in SC-β cells after transplantation and decrease in SC-α and δ cells. g, Bar graphs showing fold change differences of β-cell associated active transcription factors in SC-β cells and transplanted SC-β cells. h, Schematic showing comparison of changes associated with 6 months *in vitro* culture and 6 months *in vivo* transplants. i, Number of genes, by gene expression or ATAC promoter accessibility, or motif chromatin accessibility, downregulated or upregulated in 6 months *in vitro* and *in vivo* SC-β cells. Number of features were determined by differential gene, promoter accessibility or motif accessibility analysis comparing 2 week SC-β cells with 6 months SC-β cells *in vitro* or 6 months SC-β cells *in vivo*. j, Volcano plots showing differential motif accessibility analysis (left) and differential gene expression analysis (right) comparing 6 moths SC-β cells from *in vitro* and *in vivo* SC-islets. k, Schematic of SC-islets transfected with shRNA lentivirus for *ARID1B* gene knockdown. l, qPCR plots of SC-islets with ARID1B shRNA showing mean ± s.e.m. (n = 4) of expression of β-cell associated genes, *INS* (P = 0.0014), *DLK1* (P = 4.3 × 10^−5^), and *IAPP* (P = 3.0 × 10^−6^). Statistical significance was assessed by unpaired two-sided t-test. m, Protein quantification plot showing mean ± s.e.m. (n = 4) of human insulin content (P = 7.6 × 10^−4^) by ELISA (left), and proinsulin/insulin ratio (P = 3.6 × 10^−4^) by ELISA (right). Statistical significance was assessed by unpaired two-sided t-test. n, Volcano plot from single-cell multiomics comparing motif chromatin accessibility of SC-β cells from control and *ARID1B* shRNA condition. SC, stem cell derived; EC, enterochromaffin cells; ELISA, enzyme-linked immunosorbent assay.

To further understand the similarities and differences in endocrine cell identity between SC-islets and primary islets, we performed differential gene expression and motif chromatin accessibility analyses on the β, α, and δ-cell populations (Fig. 4g, Extended Data Fig. 5c, Supplementary Table 8). The mRNA data from the multiomic analysis yielded results consistent with previous RNA-only studies, demonstrating SC-β cells had lower expression of *IAPP, INS, UCN3, G6PC2*, and *CHGB* than primary β-cells ^9, 13, 40, 43, 45^. We also noted higher expression of *ARID1B* in SC-β cells, which is a chromatin regulator reported to play a role in identity of non-pancreatic tissues ^46^. Surprisingly, *ONECUT2* had increased mRNA expression and DNA-binding motif chromatin accessibility in SC-β cells compared to primary β-cells, despite this being a gene whose increased expression is associated with adult human β-cells compared to juvenile β-cells ^30^. The chromatin accessibility data demonstrated that SC-β cells had more enriched motifs that are associated with off-target or progenitor cell states compared to primary β-cells, such as CDX2, LMX1A, SOX9, and SOX2. Primary β-cells had enriched motifs relating to genes in the FOS/JUN family, whose gene expression is associated with improved function of SC-β cells after transplantation ^13^. MAFA surprisingly had similar chromatin accessibility of the associated DNA-binding motif between SC-β and primary β-cells, even though SC-β cells have relatively low MAFA mRNA expression and lower chromatin accessibility to the MAFA gene itself (Fig. 4f and 4g). This suggests that the role of MAFA in β-cell identity may be regulated by the chromatin state of the associated gene and not accessibility of the transcription factor to its motif. Taken together, these data illustrate that key differences exist in both mRNA expression and motif chromatin accessibility in SC-β cells compared to primary β-cells. The motif differences in particular demonstrate that SC-β cells have increased hallmarks of off-target cell types as well as a lack of key adult primary β-cell identity signatures, while primary β-cells have a more restricted chromatin state.

Earlier in Figures 1 and 3, we defined a list of the top 10 most relatively active transcription factors for each major endocrine cell for SC-islets and primary islets based on mRNA expression and motif accessibility. To better understand the differences and similarities between β, α, and δ-cells from SC-islets and primary islets, we examined the relative activity of these transcription factors in each cell type from both origins (Fig. 4h and Extended Data Fig. 5d). For the SC-β cell and primary β-cell lists, many of the transcription factors had similar mRNA expression and motif accessibility. Some notable exceptions exist, including enriched RFX1 motif accessibility, reduced *MAFA* RNA expression, and reduced NFE2L1 and BACH2 motif accessibility in SC-β cells compared to primary β-cells. To delineate the largest differences in transcription factor activity, we defined the top 10 most differentially active transcription factors in SC-β cells compared to primary β-cells. Finally, we also applied the active transcription factor list from SC-EC cells in Figure 1 to these populations. Interestingly, many of these transcription factors are enriched in SC-β cells compared to primary β-cells, further supporting the notion that SC-β cells retain some ambiguity in their cell identity due to either incorrect or incomplete cell fate specification. Taken together, these analyses highlight that while the cell types found within SC-islets generally correspond to their *in vivo* counterparts, fundamental differences still exist in cell identity in terms of mRNA expression, chromatin accessibility of gene-encoding regions, and transcription factor DNA-binding motifs

### Extended time *in vivo* but not *in vitro* improves chromatin and transcriptional SC-β cell signatures

We and others have previously demonstrated that SC-β cells can acquire improved phenotypes with extended time *in vitro* or after transplantation ^4, 9, 14, 24, 47-49^. To better characterize the chromatin and transcriptional changes that occur to cells within SC-islets over time under these two conditions, we performed multiomic sequencing on SC-islets that underwent extended culture *in vitro* or were transplanted into mice. First, stage 6 *in vitro* cultured SC-islets were sequenced at 2 weeks, 3 weeks, 4 weeks, 6 months, and 12 months (Fig. 5a, Extended Data Fig. 6a). All key SC-islet endocrine cell types (SC-β, SC-α, SC-δ, and SC-EC cells) were identified in the integrated datasets (Extended Data Fig. 6b). Interestingly, while SC-β cells displayed increased *INS* transcript with time, insulin secretion improved only until week 4 (Extended Data Fig. 6c-d). SC-α, SC-δ, and SC-EC cells demonstrated incremental increases in GCG, SST, and TPH1 expression, respectively, until week 4 but declined by month 6 (Extended Data Fig. 6c).

Similarly, we found drastic reductions in the SC-EC population at month 6 and 12 and a decrease in SC-α numbers by month 12 (Extended Data Fig. 6e). Pearson correlation analysis using RNA expression indicated that SC-EC cells began to lose their identity at month 6, while SC-α cells lost their defining signatures at month 12 (Extended Data Fig. 6f). Interestingly, Pearson correlation using the ATAC information revealed that there were drastic shifts in chromatin accessibility at month 6 and month 12 for all endocrine cell types (Extended Data Fig. 6g).

Specifically in SC-β cells, expression of many β-cell transcription factors and their associated motifs were upregulated at week 4 of *in vitro* culture, but most of these motifs and many transcripts decreased in the long term (month 6 and month 12) (Fig. 5b -d). Expression of other β-cell genes and the accessibility to their respective promoters, including *INS, IAPP*, and *UCN3*, increased over time, while the expression and promoter accessibility of genes associated with non-endocrine cells, such as the SC-EC genes LMX1A and TPH1, diminished in long term culture (Fig. 1b, d, Supplementary Table 9). Similarly, ATAC peaks demonstrated accessibility around the *INS* genomic region increased in SC-β cells over time but diminished in SC-α and SC-δ cells (Fig. 5c). Collectively, multiomic analysis of *in vitro* cultured SC-islets revealed that while the transcription of some important β-cell genes increased over time, the chromatin state of these cells became much more restricted in long-term culture. While this restriction helped to decrease off-target populations, many crucial β-cell transcription factors were also downregulated, possibly leading to the observed decrease in function.

In parallel, we also performed multiomic sequencing of SC-islets transplanted for 6 months under the kidney capsule of immunocompromised NOD.Cg-*Prkdc*^*scid*^ *Il*2*rg*^*tm1wjl*^/SzJ (NSG) mice (Fig. 5e, Extended Data Fig. 7a and b). Gene expression and ATAC promoter accessibility information demonstrated that endocrine identities became even more distinct than before transplantation (Extended Data Fig. 7c, d and e). Motif accessibility was also well defined for each cell type, with transplanted SC-β and SC-EC cells no longer displaying shared open motifs (Extended Data Fig. 7f). In particular, PDX1 was enriched in transplanted SC-β cells rather than in both SC-β and SC-EC cells as it had been before transplantation (Extended Data Fig. 7g). Comparison of these 6-month transplanted SC-islets to those before transplantation revealed that transplanted SC-islets acquired gene expression and chromatin accessibility signatures that were more similar to their primary cell counterparts, including increased *INS, MAFA*, and *IAPP* expression in SC-β cells (Extended Data Fig. 8a, b and c, Supplementary Table 10). These improvements in cell identity were also reflected in the chromatin accessibility around the *INS* gene, which exhibited diminished peak signals in non-β cell populations (Fig. 5f).

Gene set enrichment analysis illustrated that gene sets associated with β-cell identity were upregulated in transplanted SC-β cells, whereas gene sets relating to immature and off-target populations were higher in pre-transplanted SC-β cells (Extended Data Fig. 8d). Furthermore, the motif accessibility for many β-cell associated transcription factors were enriched in transplanted SC-islets (Fig. 5g). Collectively, transplanted SC-β cells were much more similar to primary β-cells than before transplantation in terms of gene expression, promoter accessibility, and motif accessibility, highlighting that transplantation improves both the transcriptional and chromatin landscape of SC-β cells (Extended Data Fig. 8f).

We then integrated the datasets of 2-week *in vitro* SC-islets, 6-month *in vitro* SC-islets, and 6-month transplanted SC-islets and found that transplanted SC-β cells had the highest enrichment of β-cell associated genes by both gene expression and promoter accessibility, such as MAFA, G6PC2, UCN3, and IAPP (Fig. 5h, Extended Data Fig. 9a and b). SC-EC cells also retained expression of identity genes when transplanted, in contrast to during 6-month *in vitro* culture. Overall, transplanted SC-islet cells displayed greater enrichment of genes and motifs for transcription factors associated with their cell types (Extended Data Fig. 9c). In particular, transplanted SC-β cells exhibited many upregulated and downregulated genes and motifs compared to their 6-month *in vitro* counterparts, allowing us to compile a list of markers relevant for β-cell identity (Fig. 5i-j, Extended Data Fig. 9d, Supplementary Table 11). In summary, extended *in vitro* culture tends to generally close chromatin regions, even some associated with β-cells. In contrast, extended time *in vivo* tends to further open chromatin regions associated with each cell’s identity while restricting regions for other lineages, allowing transplanted SC-β cells to develop a chromatin and transcriptional landscape more similar to primary β-cells.

During our SC-islet multiomic analysis, the chromatin regulator *ARID1B* emerged several times as being differentially expressed in β-cells (Fig. 4g, 5d, Extended Data Fig. 8c, 10a) and has also been reported to play a role in the identity of non-pancreatic tissues ^46^. To study its role in SC-β cell identity, we utilized short hairpin RNAs (shRNAs) targeting *ARID1B* transcript to reduce its expression in fully differentiated SC-islets (Fig. 5k and Extended Data Fig. 10b and c). *ARID1B* knockdown resulted in upregulation of β-cell identity markers, including *INS, IAPP, DLK1* (Fig 5l, Extended Data Fig. 10c). This improvement in gene expression was also associated with elevated insulin protein content, reduced pro-insulin/insulin ratio, increased amylin (IAPP) expression, and increased SC-β cell yields (Fig. 5l and m, and Extended Data Fig. 10d and e*). ARID1B* knockdown, however, did not increase either glucagon (GCG) or somatostatin (SST) protein content (Extended Data Fig. 10f). ARID1B also helped improve insulin secretion in the presence of insulin secretagogues, such as exendin 4 (Extended Data Fig. 10g). Single-cell multiomic sequencing of SC-islets with *ARID1B* knockdown (Extended Data Fig. 10h) revealed increased gene expression and motif accessibility for signatures enriched in mature human β-cells and transplanted SC-β cells, including increased chromatin accessibility around the *IAPP* genomic region (Fig 5n, Extended Data Fig. 10i and j, Supplementary Table 12). Taken together, these data demonstrate that the chromatin landscape plays a significant role in islet cell identity. Furthermore, this approach exemplifies how single-cell multiomics can be used to identify important chromatin and transcriptional deficiencies within particular cell populations that can be targeted to improve directed differentiation to specific cell types.

## Discussion

SC-β cells generated from established differentiation protocols possess many features of primary human β-cells, including the expression of β-cell specific markers, glucose responsive insulin secretion, and the ability to reverse severe diabetes in animal models ^7, 9, 12, 14, 16^. Thus, these cells have the potential to provide a functional cure for human patients with type 1 diabetes ^50^. These *in vitro*-generated cells are recognized to not have exactly the same transcriptional and functional features as their *in vivo* counterparts. An improved understanding of the deficiencies in lineage specification during differentiation to SC-islet cell types could help prevent unintended generation of non-endocrine cell types, improve endocrine cell identity, and ultimately increase SC-β cell function. By using a single-cell multiomic approach of integrating mRNA transcriptional data and ATAC chromatin accessibility information for each individual cell, we were able to generate more robust definitions of the cell types found within SC-islets than using either data set in isolation. In particular, we provide novel lists of genes and transcription factor binding motifs that emerged from this integrated analysis as being important to the specification of the different cell types found within SC-islets. Importantly, these data provide a valuable resource to identify particular deficiencies in the chromatin and transcriptional landscape of SC-islet cell types. Furthermore, we demonstrate how the differentially expressed genes and accessible DNA-binding motifs identified within this single-cell multiomic data set can be targeted to alter directed differentiation to SC-islet cell types.

Our multiomic analysis identified several interesting observations about the dynamics of gene expression and chromatin state of SC-islet cell types over time and in relation to primary human islets. In general, SC-islet cell types were less distinct from each other by chromatin accessibility compared to their primary cell counterparts. This difference was driven mainly by continued open chromatin accessibility of genes expressed by progenitor cell types, such as *NEUROG3* ^*51*^ and *GP2* ^52^, or alternative cell fates that were closed in primary cells. SC-islet cell types developed more distinguished transcriptional and chromatin accessibility signatures that matched their respective cell identities after transplantation into mice ^13, 14^, while extended *in vitro* culture broadly restricted access to chromatin regions, including those associated with a β-cell identity. The signals that the cells are receiving from the *in vivo* environment that are missing in *in vitro* culture are unknown. Furthermore, the mechanism behind this difference in cellular identity remains unclear, but similar differences may also be present within *in vitro* differentiation systems for other cell types where immaturity is commonly observed ^53, 54^.

A prior study discovered that enterochromaffin-like cells are generated during SC-islet differentiations *in vitro* and suggested that SC-EC and SC-β cells are distinct populations that arise from the same pancreatic progenitor population ^9^. By obtaining greater resolution of cell identity by simultaneously analyzing both the transcriptional and chromatin state of each cell rather than mRNA alone, our data suggests that SC-EC and SC-β cells are not completely exclusive cell populations. Instead, they comprise a continuum of cell types that contain varying degrees of both enterochromaffin and β-cell features. Interestingly, increased DNA binding motif accessibility for the chromatin regulator CTCF was highly correlative with increased SC-EC cell features. *CTCF*-overexpression blocked differentiation to pancreatic endocrine and further redirected cell fate selection toward the enteroendocrine lineage, supporting the notion that chromatin accessibility is a major regulator of the fate decision between SC-EC and SC-β cells along this gradient. As SC-EC cells may be detrimental to SC-β cell function ^9^, or at the least is unlikely to be providing a benefit for SC-islet in diabetes cell replacement therapy, understanding how to reduce or eliminate this off-target cell population is a priority for the field. In addition, the apparent plasticity of identity in SC-islets also opens up the possible of cell fate switching like has been done with primary cells ^55, 56^.

Study of SC-islets is largely motivated by success of isolated primary islet transplantation into patients with Type 1 diabetes ^57-59^. Several areas of investigation are being pursued in the field for which multiomic assessment could be beneficial. Since the early reports of generating SC-islets and other similar pancreatic endocrine ^5, 6, 60^, many teams has focused on changing the composition of the media or how the cells are cultured to improve SC-islet function ^4, 7, 8, 14, 61, 62^. Controlled assembly of the three-dimensional SC-islet aggregates and circadian entrapment have also further improved function of SC-islets ^4, 8, 24^. Proliferation of early endoderm and pancreatic progenitors have also been achieved to expand the number of cells available for continued differentiation to SC-islets and potentially improve them ^63-66^. Increased efficacy of generation or purification of progenitors have also been pursued to generate SC-islets ^10, 52, 67, 68^. Furthermore, genetic engineering to lessen immune recognition ^69^ of SC-islets in order to transplant the cells without immunosuppression has been reported ^62, 70^. SC-islets have been generated from patients with diabetes for study of diabetes pathology and consideration as an autologous cell source of diabetes regenerative medicine ^12, 71-76^. Finally, many materials are being studied to provide immune protection or otherwise benefit islets or SC-islets after transplantation ^77-84^.

Our data demonstrating that chromatin accessibility differs drastically between cell types as well as within a given cell type from different origins, such as *in vitro* differentiation compared to isolation from donors, suggesting that regulators of chromatin may play a substantial role during the generation and identity of islet cell types. Our modulations of the chromatin regulators CTCF and ARID1B to alter SC-β cell identity, highlight how better control of the chromatin landscape could lead to the development of SC-islet differentiation protocols with improved function and defined cellular composition. Here, we provide a comprehensive indexing of single-cell transcriptional state and chromatin accessibility in SC-islets to help facilitate further development of these differentiation protocols, as both aspects of islet cell identity will likely need to be targeted to enhance SC-islet differentiation strategies.

## Methods

### Cell culture and differentiation

The HUES8 hESC line was provided by Douglas Melton (Harvard University)^5^. The H1 hESC line was provided by Lindy Barrett (Broad Institute) with permission from WiCell containing doxycycline-inducible dCas9-VPR transgene in the AAVS1 locus (CRISPRa system)^39^. hESC work was approved by the Washington University Embryonic Stem Cell Research Oversight Committee (approval no. 15-002) with appropriate conditions and consent. mTeSR1 (StemCell Technologies; 05850) was used for the culture of undifferentiated cell propagation. All cell culture was maintained in a humidified incubator at 5% CO_2_ and 37°C. Cells were passaged every 4 days by washing cell with PBS and incubating with 0.2 mL TrypLE/cm^2^ (Gibco; 12-604-013) for 10 min or less at 37°C. Cells were collected into a 15-mL conical containing equal volume mTeSR1 supplemented with 10 µM Y-27632 (Pepro Tech; 129382310MG). Cells were counted on Vi-Cell XR (Beckman Coulter) and spun at 300g for 3 min at RT. The supernatant was aspirated, and cells were seeded at a density of 0.8×10^5^/cm^2^ for propagation onto a Matrigel (Corning; 356230) coated plate in mTeSR1 supplemented with 10µM Y-27632 (Pepro Tech; 129382310MG). After 24 hr, media was replaced daily with mTeSR1 alone. SC-islet differentiation was performed as described previously^7, 29^. Briefly, 24 hr after seeding hESCs at a density of 6.3×10^5^ cells/cm^2^, daily mediums with added growth factors were started to initiate the differentiation process. The base media formulation and added growth factors can be found in Supplementary Table 1. After 7 days in stage 6, cells were placed on an Orbishaker (Benchmark) at 100 RPM and maintained in ESFM media. All *in vitro* data in the paper is from 14 days in stage 6 unless otherwise noted. Human islets were acquired from Prodo Laboratories for comparison in some experiments.

### Single-nuclei sample preparation and sequencing

Cells collected were processed and delivered to the McDonnell Genome Institute at Washington University for library preparation and sequencing. Single-cell suspension samples were processed into nuclei according to 10X Multiome ATAC + Gene Expression (GEX) protocol (CGOOO338). Briefly, cell samples were collected and washed with PBS (with 0.04% BSA), lysed with chilled Lysis Buffer for 4 min, washed 3 times with wash buffer, and resuspended with 10x nuclei buffer at 3000-5000 nuclei/μl. Nuclei samples were subsequently processed using the Chromium 10x genomics instrument, with a target cell number of 7000-10000. The 10x Single Cell Multiome ATAC + Gene Expression v1 kit was used according to the manufacturer’s instructions for library preparations. Sequencing of the library was performed using the NovaSeq 6000 System (Illumina).

### Multiomics sequencing analysis workflow

Multiomic sequenced files were processed for demultiplexing and analyzed using Cell Ranger ARC v2.0. Genes were mapped and referenced using GRCh38. RStudio with R version 4.0.3 was used to perform subsequent analyses. Datasets were further analyzed using Seurat 4.01^85^ and Signac 1.3.0^86^. For ATAC data, genomic positions were mapped and annotated with EnsDb.Hsapeins.v86 and hg38. Low quality cells including doublets, dead cells, and poor sequencing depth cells, were filtered by removing cells with low RNA counts (nCount_RNA < 1000) and low ATAC counts (nCount_ATAC < 1000); high RNA counts (ranging > 40000 – 50000) and high ATAC counts (ranging > 40000 – 50000); nucleosome signal > 1.25, and TSS enrichment < 2. In addition, mouse cells were removed by removing cells showing high expression of kidney marker *TTC36*^13^ for transplanted SC-islet samples. Additional details for filtering cells can be found in Supplementary Table 2 and Supplementary Figure 1. Gene expression data and ATAC data were processed using SCT transform for the former and RunTFIDF and RunSVD for the latter. Dataset integration was performed by anchoring using SCT normalized data and integrative gene expression and ATAC dimension reductions were carried out using PCA and LSI. Clusters were identified using differential gene expression analysis or expression of known marker genes. Promoter accessibilities were determined from ATAC information using GeneActivity by considering 2000 base pairs upstream of the transcription start site. ATAC peaks were called using MACS2 and linked using LinkPeaks and Cicero to determine cis-regulatory elements^86, 87^. Motif activity was computed by using the chromVAR package^88^ and JASPAR version 2020 database^89^. Trajectory analysis was conducted using Monocle3 package^90^.

### Transduction of *CTCF* gRNA in CRISPRa line

CRISPRa genetic engineering of the H1 dCas9-VPR line^39^ was mediated using custom guide RNAs (gRNAs) ordered from MilliporeSigma. *CTCF* targeting gRNAs for CRISPRa sequences are provided in Supplementary Table 1. gRNAs were resuspended to a final concentration of 100uM in water. Primers were phosphorylated and ligated together by adding T4 ligation buffer and T4 Polynucleotide kinase (PNK) enzyme (NEB; B0202A and M0201S) and running on a thermocycler under the following conditions: 37 °C for 30 min; 95 °C for 5 min; and ramp down to 25 °C at 5 °C/min. Oligos were then diluted with 90 µl of ultrapure water. These oligos were then inserted into the sgRNA library backbone (Lenti sgRNA(MS2)_puro) using the golden gate reaction. This is achieved by adding a 25ng ul plasmid backbone to a master mix of Rapid Ligase Buffer 2X (Enzymatics: B1010L), Fast Digest Esp31 (Thermo: FD0454), DTT (Promega: PRP1171), BSA (NEB: B9000S), T7 DNA ligase, (Enzymatics: L6020L) and the diluted gRNA oligos in a total reaction volume of 25 ul. The Golden Gate assembly reaction is then preformed in a thermocycler under the following conditions: 15 cycles of 37 °C for 5 min, 20 °C for 5 min with final hold at 4°C. Lenti sgRNA (MS2) puromycin optimized backbone was a gift from Feng Zhang (Addgene plasmid # 73797). This final plasmid was then transfected into STBL3, following the same methods as mentioned in the previous shRNA lentivirus production section. To transfect the CRISPRa H1 dCas9-VPR stem cell line, lentiviral particles containing gRNA was added at MOI 5 with polybrene (5 µg/mL) in culture for 24 hr. At confluency, transfected stem cells were passaged and cultured with media containing puromycin (1 µg/mL) for selection. To induce CRISPRa expression, doxycycline (MilliporeSigma) was added at 1 µg/mL for 7 d during stage 5 of the differentiation protocol.

### Real-Time PCR

Cells were lysed directly with RLT buffer from the RNeasy Mini Kit (74016; Qiagen) followed by RNA extraction following the manufacturer’s instructions. cDNA was synthesized from the RNA using the High-Capacity cDNA Reverse Transcription Kit (Applied Biosystems; 129382310MG) on a T100 thermocycler (BioRad). PowerUp SYBR Green Master Mix (Applied Biosystems; A257411) was used to run samples on the Quant Studio 6 Pro (Applied Biosystems) and results were analyzed using ΔΔCt methodology. Primer sequences used in this paper are listed in Supplementary Table 1.

### Microscopy

Fluorescence images were taken on a Zeiss Cell Discoverer confocal 7 microscope. For immunocytochemistry (ICC), cells were fixed in 4% paraformaldehyde (PFA) (Electron Microscopy Science; 15714) for 30 min at RT. For staining, fixed cells were blocked in ICC solution (PBS (Fisher; MT21040CV), 0.1% Triton X (Acros Organics; 327371000), 5% donkey serum (Jackson Immunoresearch; 01700-121)) for 30 min at RT. Samples were subsequently treated with primary and secondary antibodies in ICC solution overnight at 4°C and 2 hr at RT respectively. DAPI (Invitrogen; D1306) was used for nuclear staining. Samples were incubated in DAPI for 12 min at RT, washed with ICC solution and stored in PBS until imaging. Antibody details and dilutions can be found in Supplementary Table 1. Image J was used for analysis. Quantification was performed by manual counting of cells from analyzed fluorescent images and can be found in Supplementary Figure 2.

### Flow cytometry

Cells were single-cell dispersed by washing with PBS and adding TrypLE/cm^2^ for 10 min at 37°C. Samples were washed with PBS, fixed for 30 min in 4% PFA at 4°C and washed with PBS. For staining, samples were treated with ICC solution for 45 min at RT. Primary antibodies were prepared in ICC solution and incubated on cells overnight at 4°C. Samples were washed with PBS and incubated for 2 hr with secondary antibodies in ICC at 4°C. Cells were washed twice with PBS and filtered before running on the LSR Fortessa flow cytometer (BD Bioscience). FlowJo v10.8.0 (Becton, Dickinson, and Company) was used for analysis. Gating strategy for flow cytometry analysis can be found in Supplementary Figure 3.

### Hormone content

Insulin, somatostatin, glucagon, and proinsulin were measured from differentiated SC-islets. Cells were collected, rinsed, and incubated in acid-ethanol solution for 48 hr at -20°C. Sample solutions were neutralized with 1M TRIS buffer (Millipore Sigma; T6066) for downstream processing. Measurements of insulin, somatostatin, glucagon and proinsulin were made using the following ELISA kits: Human insulin ELISA (ALPCO; 80-INSHU-E01.1), Somatostatin EIA (Phoenix Pharmaceuticals; EK-060-03), Glucagon ELISA (Crystal Chem; 81520), and human proinsulin ELISA (Mercodia; 10-1118-01). ELISAs were performed according to the manufacturer instructions. Results were normalized to cell counts performed on the Vi-Cell XR (Beckman Coulter).

### Mouse transplantations and SC-islet cell retrieval

Mice that were 7-week old, male, and with the NOD.Cg-*Prkdc*^*scid*^ *Il*2*rg*^*tm1wjl*^/SzJ (NSG) background (Jackson Laboratories; 005557) were randomly assigned to experimental groups. Mice were housed in a facility with a 12-hr light/dark cycle and were fed on chow diet. Animal studies were done in accordance with Washington University International Care and Use Committee (IACUC) guidelines (Approval 21-0240). Animal studies were performed by unblinded individuals in accordance with Washington University International Care and Use Committee (IACUC) guidelines. Mice randomly assigned to experimental groups were anesthetized using isoflurane and injected with ∼5×10^6^ SC-islets under the kidney capsule. At 6 months post-transplantation, mice were euthanized, and the kidney transplanted with SC-islets was removed. A razor blade was used to mince the kidney prior to placing it into a solution of 2 mg/mL collagenase D (Sigma; 11088858001) in RPMI (GIBCO; 1187-085). The tissue was incubated for 40 min at 37°C before diluting with PBS and mechanical disrupting with a pipette and filtering through a 70-µm strainer (Corning; 431751). The flow through was centrifuged and the remaining cell pellet was resuspended in MACS buffer (0.05% BSA in PBS). The Miltenyi mouse cell isolation kit (Miltenyi; 130-104-694; LS column, 130-042-401) was used to remove any excess mouse cells from the cell solution. The flow through of cells was collected, centrifuged, counted, and resuspended in PBS with 0.04% BSA for nuclei processing and sequencing.

### Lentiviral design, preparation, and transduction

Gene KD of *ARID1B* was performed using pLKO.1 TRC plasmids containing shRNA sequences targeting *ARID1B* and *GFP* (Control) (Supplementary Table 1). Glycerol stocks were grown, and plasmid DNA was isolated using Qiagen Mini-prep kit (Qiagen; 27115). Plasmid DNA was transfected into One Shot™ Stbl3™ Chemically Competent E. coli (Invitrogen; C737303) and spread on an agar plate. After 18 hr a single colony was selected, cultured, and DNA was extracted using the Qiagen Maxi-prep-plus kit (Qiagen; 12981). Viral particles were generated using Lenti-X 293T cells (Takara; 632180) cultured in DMEM with 10% heat inactivated fetal bovine serum (MilliporeSigma; F4135), and 0.01mM Sodium Pyruvate (Corning; 25-000-CL) in 10cm tissue culture treated plates (Falcon; 353003). Confluent Lenti-X 293T cells were transfected with 6 μg of shRNA plasmid, 4.5 μg of psPAX2 (Addgene; 12260; gift from Didier Trono), and 1.5 μg pMD2.G (Addgene; 12259; gift from Didier Trono) packaging plasmids in 600 μL of Opti-MEM (Life Technologies; 31985-070) and 48 μL of Polyethylenimine ‘Max’ MW 40,000 Da (Polysciences; 24765-2) per plate. Media was refreshed after 16 hr. Viral containing supernatant was collected at 96 hr post transfection and concentrated using Lenti-X concentrator (Takara; 631232). Collected lentivirus was tittered using Lenti-X™ GoStix™ Plus (Takara;631280). Lentiviral transduction was initiated at the first day of Stage 6 with an MOI of 5 for 24 hr.

### Statistics and reproducibility

Cell selection criteria or exclusion methods for single-cell analysis can be found in Supplementary Figure 1. For *in vitro* experiments, we performed unpaired or paired parametric t-tests (two-sided) to determine significance. All *in vitro* experiment data points presented are biological replicates. Significant values are marked based on p-values using ns > 0.05, * < 0.05, ** < 0.01, *** < 0.001, and **** < 0.0001.

## Data availability

Single-cell multiomics sequencing analysis and source data can be found in the supplementary tables. Fluorescent images for quantification can be found in Supplementary Figure 2. Individual flow cytometry data can be found in Supplementary 4. All sequencing datasets have been deposited in the Gene Expression Omnibus (GEO) – NCBI under accession number GSE199636.

## Code availability

Codes used for analyzing single-nuclei multiomic sequencing are available on https://github.com/punnaug.

## Acknowledgements

This work was funded by NIH (R01DK114233, R01DK127497), JDRF (5-CDA-2017-391-A-N), and startup funds from the Washington University School of Medicine Department of Medicine. N.J.H. was supported by a JDRF Advanced Postdoctoral Fellowship (3-APF-2020-930-A-N). M.I. was supported by Rita Levi-Montalcini Postdoctoral Fellowship in Regenerative Medicine and the NIH (T32DK007120). M.M.M. was supported by the NIH (T32GM139774). J.R.Miller was supported by a Washington University BioSURF award. D.A.V.P. was supported by the NSF Graduate Research Fellowship Program (DGE-2139839 and DGE-1745038). L.V.C. was supported by the NIH (F31DK125068). Microscopy was performed through the Washington University Center for Cellular Imaging, which is supported by the Washington University School of Medicine, the Children’s Discovery Institute (CDI-CORE-2015-505) and the Foundation for Barnes-Jewish Hospital (3770). Microscopy analysis was supported by the Washington University Diabetes Research Center (P30DK020579) and Center of Regenerative Medicine. We thank the Genome Technology Access Center at the McDonnell Genome Institute at Washington University School of Medicine for help with genomic analysis, which is supported by NIH (P30CA91842) support to the Siteman Cancer Center and by ICTS/CTSA (UL1TR002345) from the National Center for Research Resources (NCRR). We thank N. Udomkittivorakul for the graphics design, D. Melton (Harvard University) for the HUES8 cell line, and L. Barret (Broad Institute of MIT and Harvard University) for the CRISPRa VPR cell line.

## Contributions

P.A. and J.R.Millman designed all experiments and performed all *in vivo* experiments. P.A. performed all computational analysis and associated cell culture. P.A., E.M., M.M.M., M.D.S., M.I., D.A.V.P., J.R.Miller, S.E.G., and L.V.C. performed *in vitro* experiments. P.A., N.J.H., M.I., and J.R.Millman wrote the manuscript. All authors revised and approved the manuscript.

## Additional information

Supplementary Information is available for this paper. Correspondence and requests for materials should be addressed to J.R.Millman Reprints and permissions information is available at www.nature.com/reprints.

