## Extended Data for "Defining the chromatin and transcriptional landscape of stem cell-derived islets"

Washington University School of Medicine

MSC 8127-057-08

660 South Euclid Avenue

St. Louis, MO 63110

USA

[2]Department of Biomedical Engineering

Washington University in St. Louis

1 Brookings Drive

St. Louis, MO 63130

USA

*To whom correspondence should be addressed:

Jeffrey R. Millman,

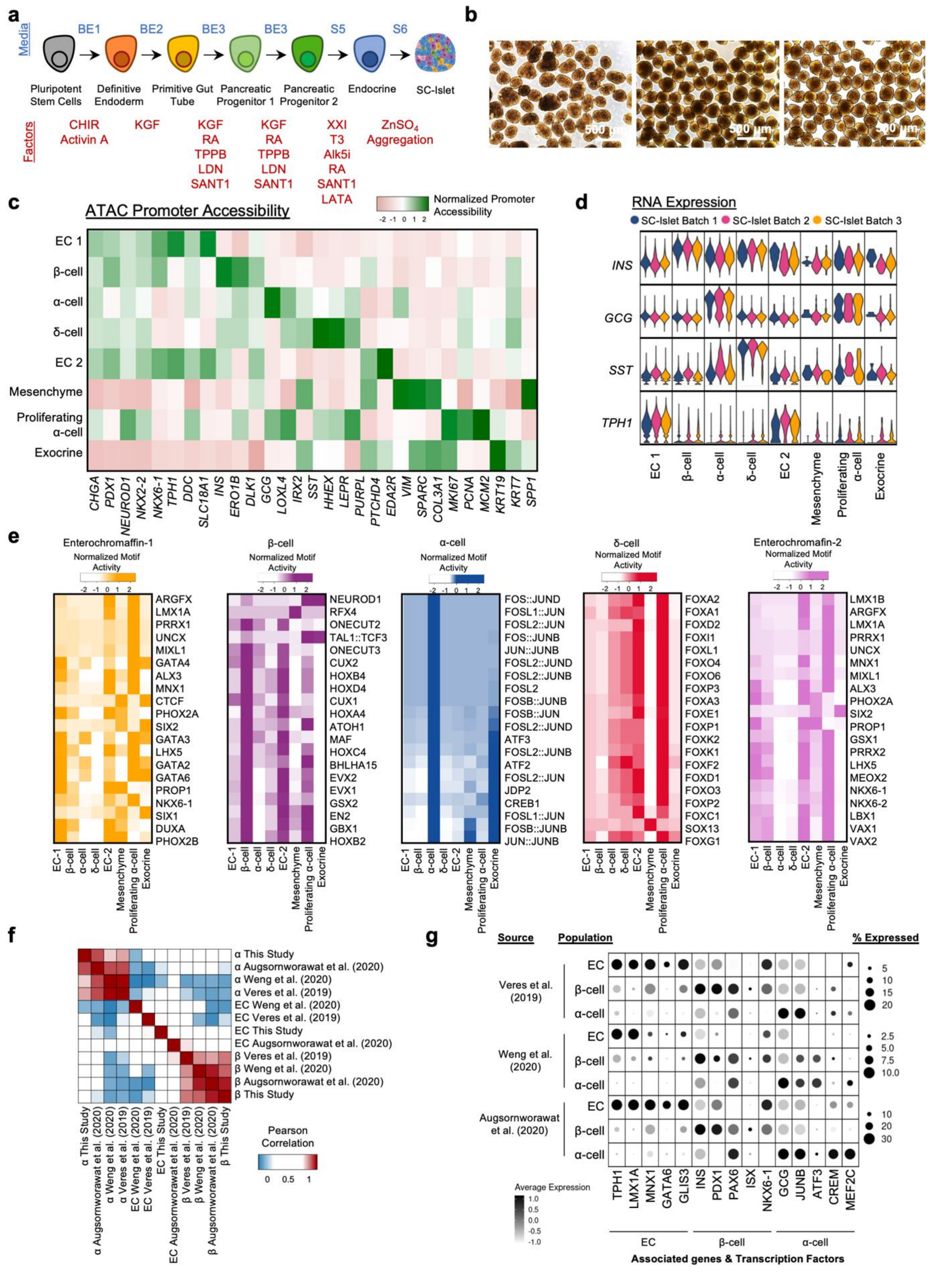


**Extended Data Fig. 1. | Single-cell Multiomic ATAC and gene expression characterization of SC-islets.** a, Schematic diagram outlining the differentiation protocol for generating SC-islets.

b, Brightfield images of differentiated SC-islets used for multiomic sequencing. c, Heatmap showing ATAC promoter accessibility of markers associated with each cell type. d, Violin plots for INS, GCG, SST, and TPH1 gene expression in SC-islet cell types from different datasets of different differentiation batches. e, Heatmaps showing the top 20 enriched motifs in SC-EC1, SC-β, SC-α, SC-δ, and SC-EC2 cells. f, Pearson correlation analysis using the top 2000 variable genes comparing correlation of SC-β, SC-α, and SC-EC in SC-islets from this study and other literatures using other differentiation protocols. SC-β, and SC-α have similar gene expression profiles across multiple datasets. g, Dotplots showing the upregulation, by gene expression, of identified active transcription factors in SC-β, SC-α, and SC-EC cells of datasets from other studies. EC, enterochromaffin cells.

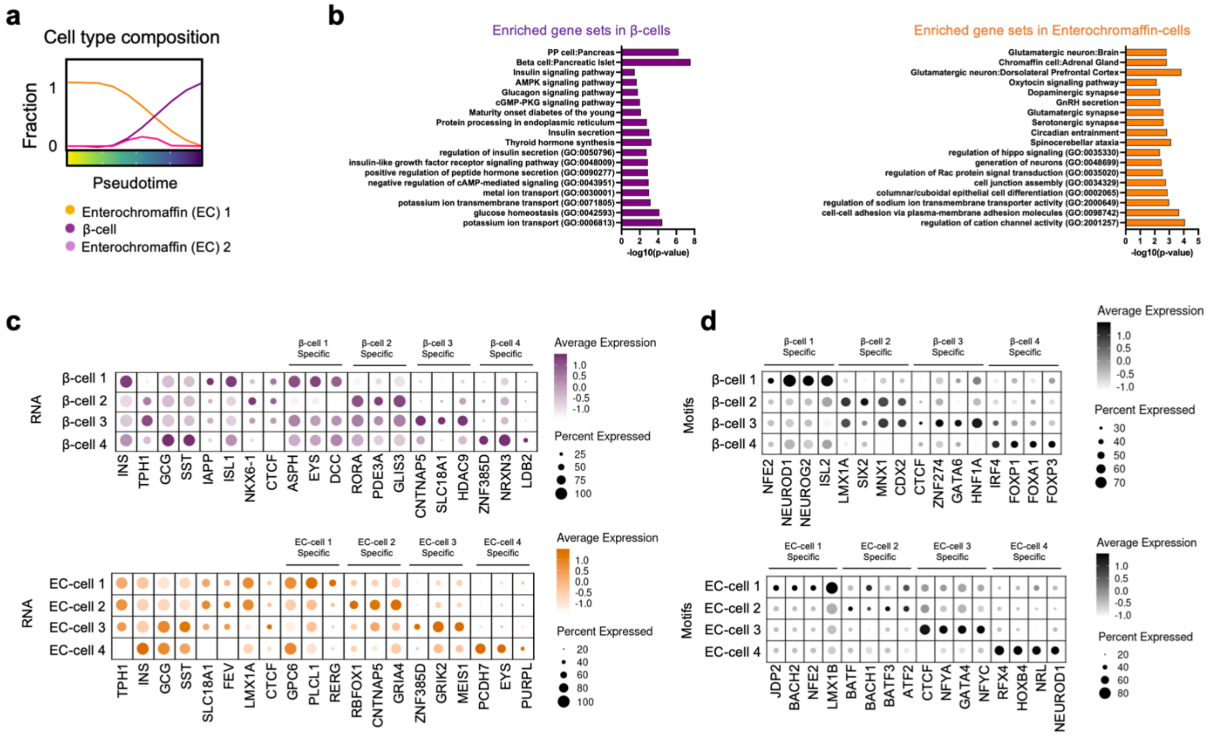


**Extended Data Fig. 2. | Delineation of SC-β and SC-EC analysis.** a, Composition of cell types represented on the trajectory analysis. b, Gene set enrichment of analysis showing enrichment of gene sets in segments of trajectory representing SC-β and SC-EC cells. c, Dotplots showing marker genes and unique genes associated with subpopulations identified from the reclustering of SC-β and SC-EC cell populations. d, Dotplots showing unique motif chromatin accessibility in in subpopulations of SC-β and SC-EC cells. EC, enterochromaffin cells.

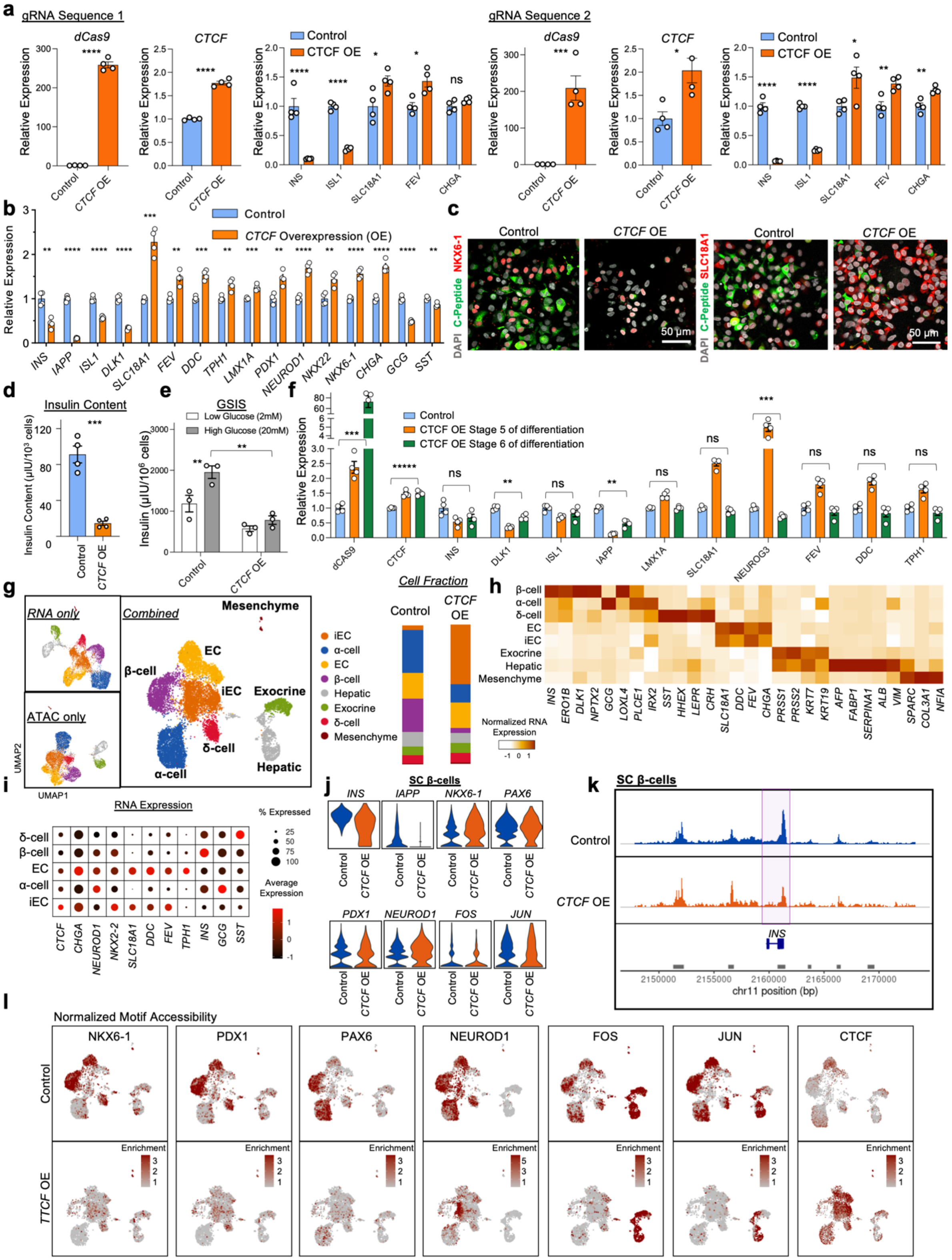


**Extended Data Fig. 3. | *CTCF* induction during differentiation using doxycycline inducible CRISPRa stem cells.** a, qPCR analysis, plotting mean ± s.e.m. (n = 4), showing upregulation of dCas9 (gRNA sequence 1, P = 4.2 x $10^{-8}$; gRNA sequence 2, P = 7.8 x $10^{-4}$), *CTCF* overexpression (gRNA sequence 1, P = 1.4 x $10^{-6}$; gRNA sequence 2, P = 0.014), and changes in expression of SC-β and SC-EC marker genes (gRNA sequence 1, *INS* (P = 5.6 x $10^{-4}$) *ISL1* (P = 1.1 x $10^{-6}$), *SLC18A1* (P = 0.034), *FEV* (P = 0.012), *CHGA* (ns, P = 0.12); gRNA sequence 2, *INS* (P = 4.4 x $10^{-6}$), *ISL1* (P = 7.8 x $10^{-8}$), *SLC18A1* (P = 0.042), *FEV* (P = 0.0069), *CHGA* (P = 0.0050), upon activation using doxycycline. b, qPCR analysis, plotting mean ± s.e.m. (n = 4), of *CTCF* overexpressed SC-islets after differentiation showing expression differences of genes associated with β cells (*INS*, P = 0.0010; *IAPP*, P = 1.5 x $10^{-7}$; *ISL1*, P = 3.0 x $10^{-5}$; *DLK1*, P = 3.7 x $10^{-6}$), EC cells (*SLC18A1*, P = 1.6 x $10^{-4}$; *FEV*, P = 0.0019; *DDC*, P = 1.2 x $10^{-4}$; *TPH1*, P = 0.0047; *LMX1A*, P = 4.6 x 10-4), pancreatic identity (*PDX1*, P = 0.0031; *NEUROD1*, P = 6.2 x $10^{-5}$; *NKX2-2*, P = 0.0047; *NKX6-1*, P = 7.5 x $10^{-5}$; *CHGA*, P = 6.8 x $10^{-5}$), and other islet hormones (*GCG*, P = 5.9 x $10^{-6}$; *SST*, P = 0.0046). c, Immunocytochemistry of differentiated SC-islets with *CTCF* overexpression showing diminished SC-β population with C-peptide (green) and NKX6-1 (red) and increased SC-EC cell with SLC18A1 (red). d, Protein quantification plot showing mean ± s.e.m. (by ELISA, n = 4) of decreased human insulin content after CTCF overexpression (P = 2.2 x $10^{-4}$). Statistical significances were assessed by unpaired two-sided t-test. e, Glucose stimulated insulin secretion assay, plotting mean ± s.e.m. (by ELISA, n = 3), of differentiated control SC-islets (P = 0.0095) and SC-islets with *CTCF* overexpression. SC-islets with *CTCF* overexpression displays lower insulin secretion at high glucose stimulation when compared to control (P = 0.0032). Statistical significances were assessed using paired two-sided t-test and unpaired two-sided t-test respectively. f, qPCR analysis, plotting mean ± s.e.m. (n = 4), of differentiated SC-islets comparing CTCF overexpression during endocrine induction (Stage 5 of protocol) or at the end of differentiation (Stage 6 of protocol). CTCF overexpression at the end of the differentiation exhibits some decreased gene expression of β cell genes, but no significant increase in EC cell associated genes. (*dCAS9*, P = 6.13 x $10^{-4}$; *CTCF*, P = 2.0 x $10^{-5}$; *INS*, P = 0.10; *DLK1*, P = 0.0023; *ISL1*, P = 0.18; *IAPP*, P = 3.3 x $10^{-4}$; *LMX1A*, P = 0.97; *SLC18A1*, P = 0.12; *NEUROG3*, P = 9.3 x $10^{-4}$; *FEV*, P = 0.30; *DDC*, P = 0.27; *TPH1*, P = 0.21) g, UMAPs showing identified cell types in SC-islets induced with *CTCF* overexpression using both or either ATAC or gene (RNA) information (left). Composition of cell types represented in the UMAP (right). h, Heatmap showing normalized gene expression of genes associated with identified populations in the *CTCF* CRISPRa experiment. i, Dot plots highlighting the gene expression of *CTCF* and endocrine-associated genes in the SC-β, SC-α, SC-δ, SC-EC, and SC-iEC cell populations. j, Violin plot of β-cell associated gene expressions of CTCF overexpression in the SC-β cell population. k, Chromatin accessibility around the *INS* genomic region of SC-β cells, showing reduced peaks in the CTCF overexpression condition. l, Feature plots showing decreased motif accessibility of β cell associated motifs and increased accessibility to CTCF motif in endocrine populations with CTCF overexpression. EC, enterochromaffin cells; ns, not significant.

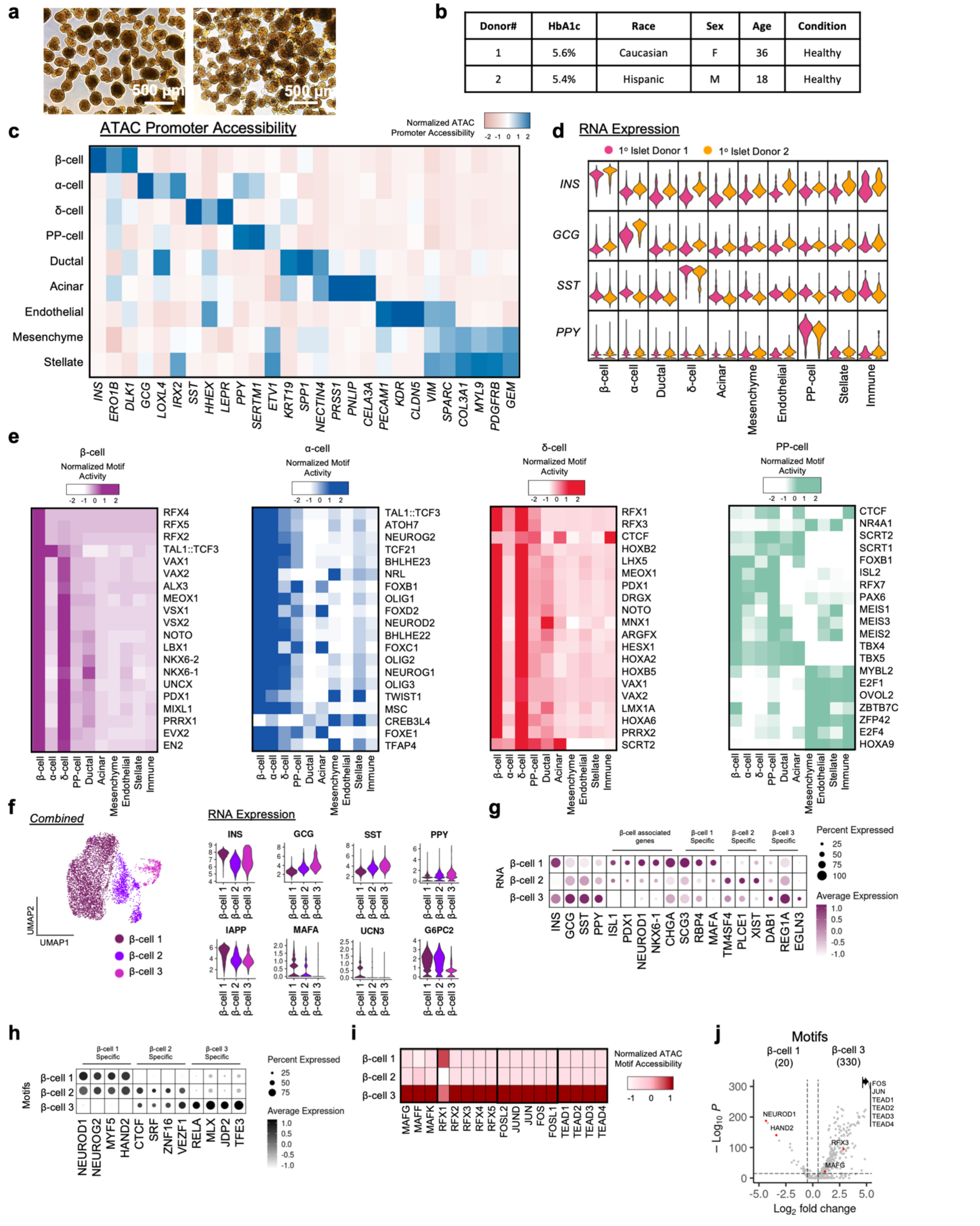


| Donor# | HbA1c | Race | Sex | Age | Condition |
|---|---|---|---|---|---|
| 1 | 5.6% | Caucasian | F | 36 | Healthy |
| 2 | 5.4% | Hispanic | M | 18 | Healthy |

**Extended Data Fig. 4. | Single-cell Multiomic ATAC and gene expression characterization of primary huma islets.** a, Brightfield images of primary islets used for multiomic sequencing.

b, Primary islet donor information. c, Heatmap showing ATAC promoter accessibility of markers associated with each cell type. d, Violin plots for INS, GCG, SST, and PPY gene expression in primary islet cell types from different donors. e, Heatmaps showing the top 20 enriched motifs in primary β, α, δ, and PP cells. f, Re-clustering analysis of primary β cells highlighting heterogeneity by variations in mature and polyhormonal gene expressions. g, Dotplots showing marker genes and unique genes associated with subpopulations identified from the re-clustering of primary β cells. h, Dotplots showing unique motif chromatin accessibility in in subpopulations primary β cells. i, Chromatin accessibility of transcription factor binding sites of MAF, RFX, FOS/JUN, and TEAD motif family in primary β cell subpopulations. j, Volcano plots comparing motif chromatin accessibility between INS high β-cell 1 and INS low β-cell 2 subpopulation. PP, Pancreatic progenitors.

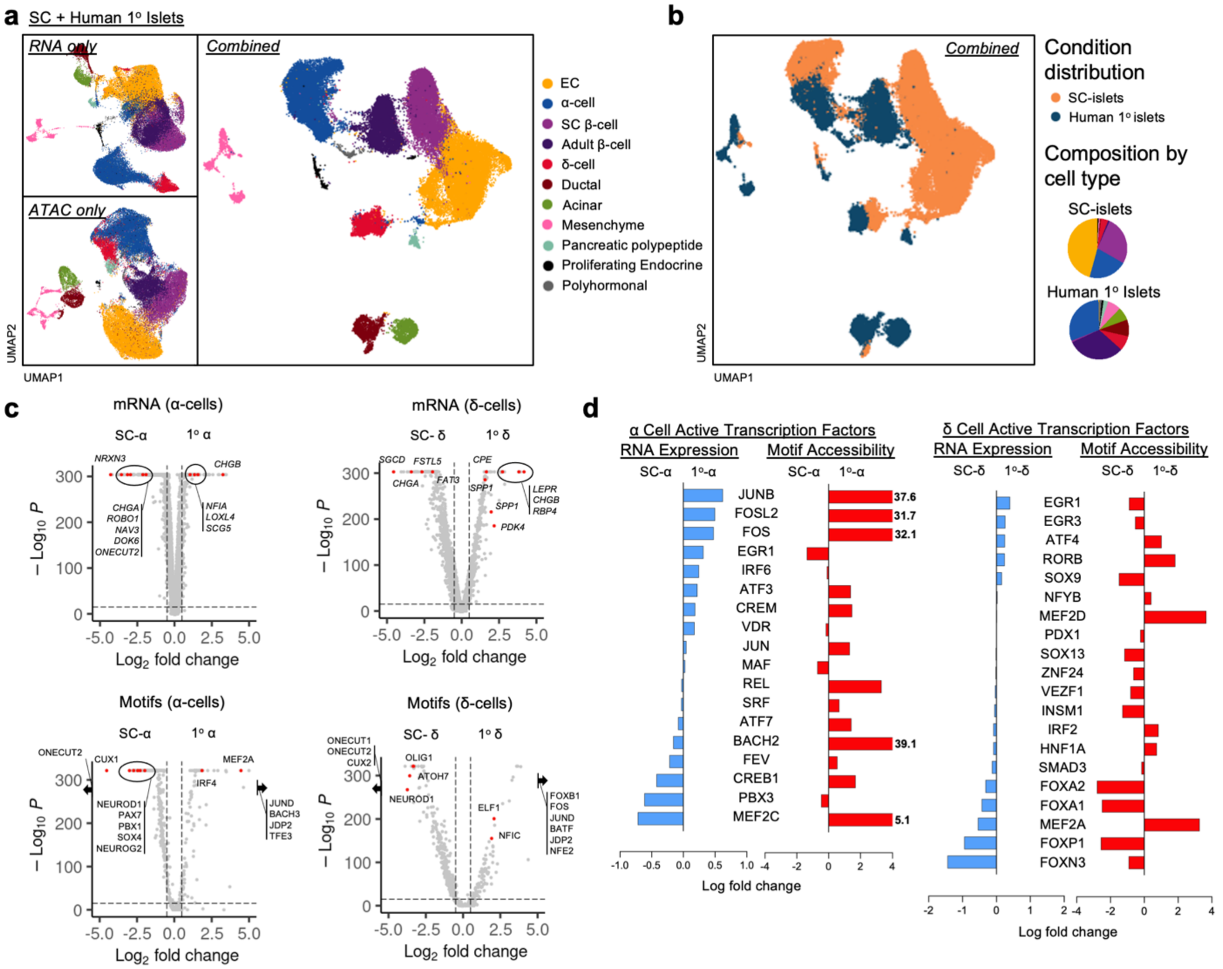


**Extended Data Fig. 5. | Multiomic sequencing comparison of SC-islets and primary human islets.** a, Integrative UMAP showing cells from SC-islets and primary islets. b, Integrative UMAP plotted by SC-islet or primary islet condition. Pie charts show composition information of cell types identified. c, Differential gene expression (top) and motif chromatin accessibility analysis (right) for α-cells (left) and δ-cells (right). d, Bar graphs showing fold change comparing SC and primary α and δ cell populations, showing gene expression and motif chromatin accessibility of identified transcription factors associated with the respective cell types. EC, enterochromaffin cells.

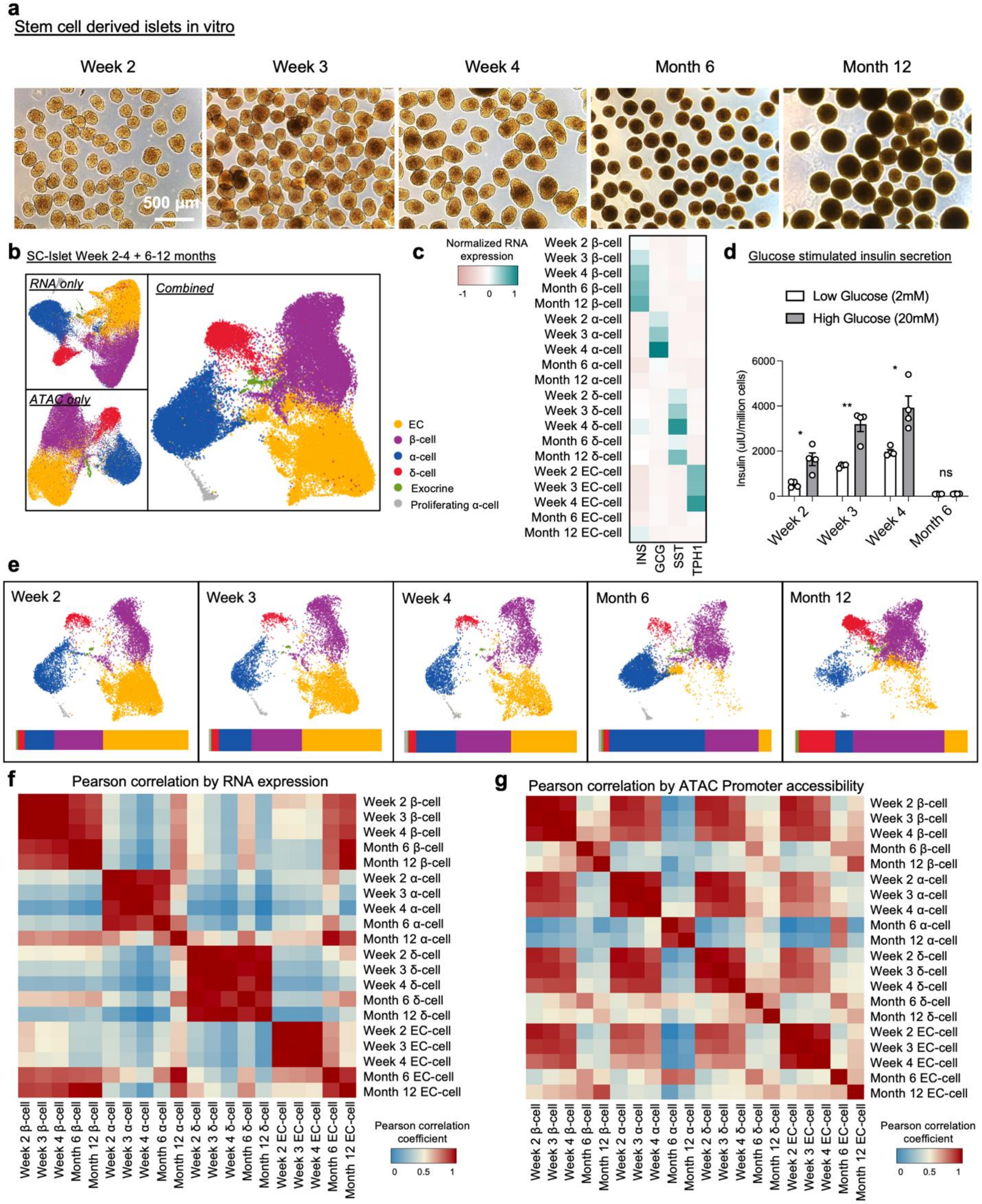


**Extended Data Fig. 6. | Time course characterization and analysis of long-term in vitro SC-islets.** a, Bright field images of SC-islets from week 2, week 3, week 4, month 6, and month 12

of in vitro culture. b, Integrative UMAP of in vitro SC-islet cells including all time points. c, Heatmap showing increase, or decrease, of marker gene expression in SC-islet cells through out time. d, Glucose stimulated insulin secretion assay, plotting mean ± s.e.m. (by ELISA, n = 4), of SC-islets cultured in vitro at week 2 (P = 0.020), week 3 (P = 0.0080), week 4 (P = 0.046), and month 6 (P = 0.27) time point. e, UMAP and bar plots showing composition information of cell types from integrated time course datasets. f and g, Pearson correlation to compare all timepoints, of cell types, (f) using top 2000 most variably expressed genes (mRNA), and (g) using top 2000 most variably accessible promoters from ATAC. EC, enterochromaffin; ns, not significant.

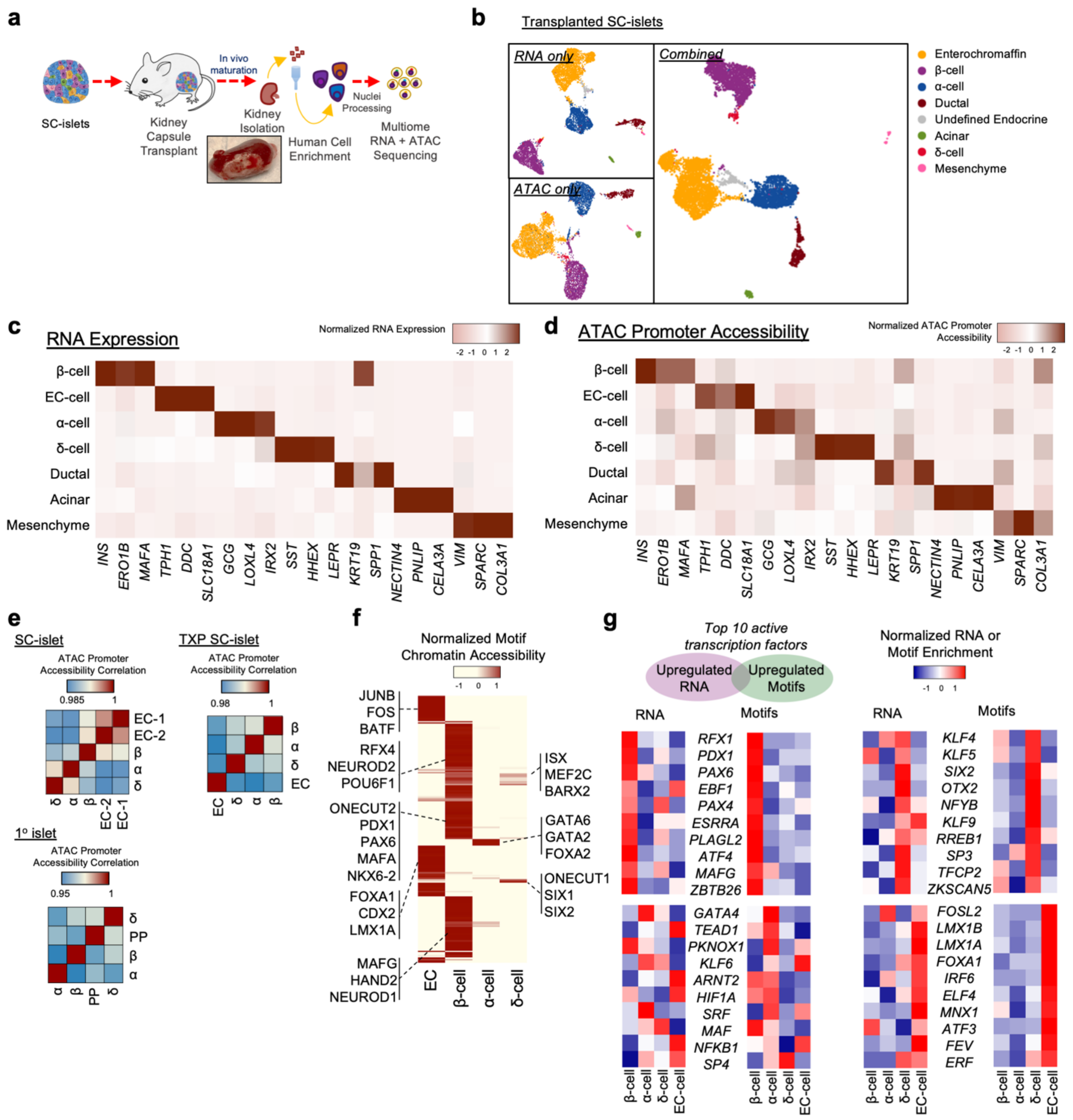


**Extended Data Fig. 7. | Single-cell Multiomic ATAC and gene expression characterization of transplanted SC-islets.** a, Schematics of SC-islet transplantation and retrieval of graft after 6 months in vivo. b, UMAP and identification of transplanted SC-islet cells using gene expression and chromatin information. c and d, Heatmap showing gene expression (c) and ATAC promoter accessibility (d) of markers associated with each cell type. e, Pearson correlation analysis using top 2000 most variable ATAC promoter accessibility of key endocrine populations from SC-islets, transplanted SC-islets, and primary human islets. This analysis highlights the distinctiveness of chromatin identity acquired in cell types from SC-islets after transplantation. f, Heatmap showing the top 200 variable motifs within endocrine cell populations and highlighting motif markers for each in vivo cell type. f, Heatmaps highlighting gene expression and ATAC motif accessibility of top 10 active transcription factors co-enriched with both features in transplanted SC-β, SC-α, SC-δ, and SC-EC cells.

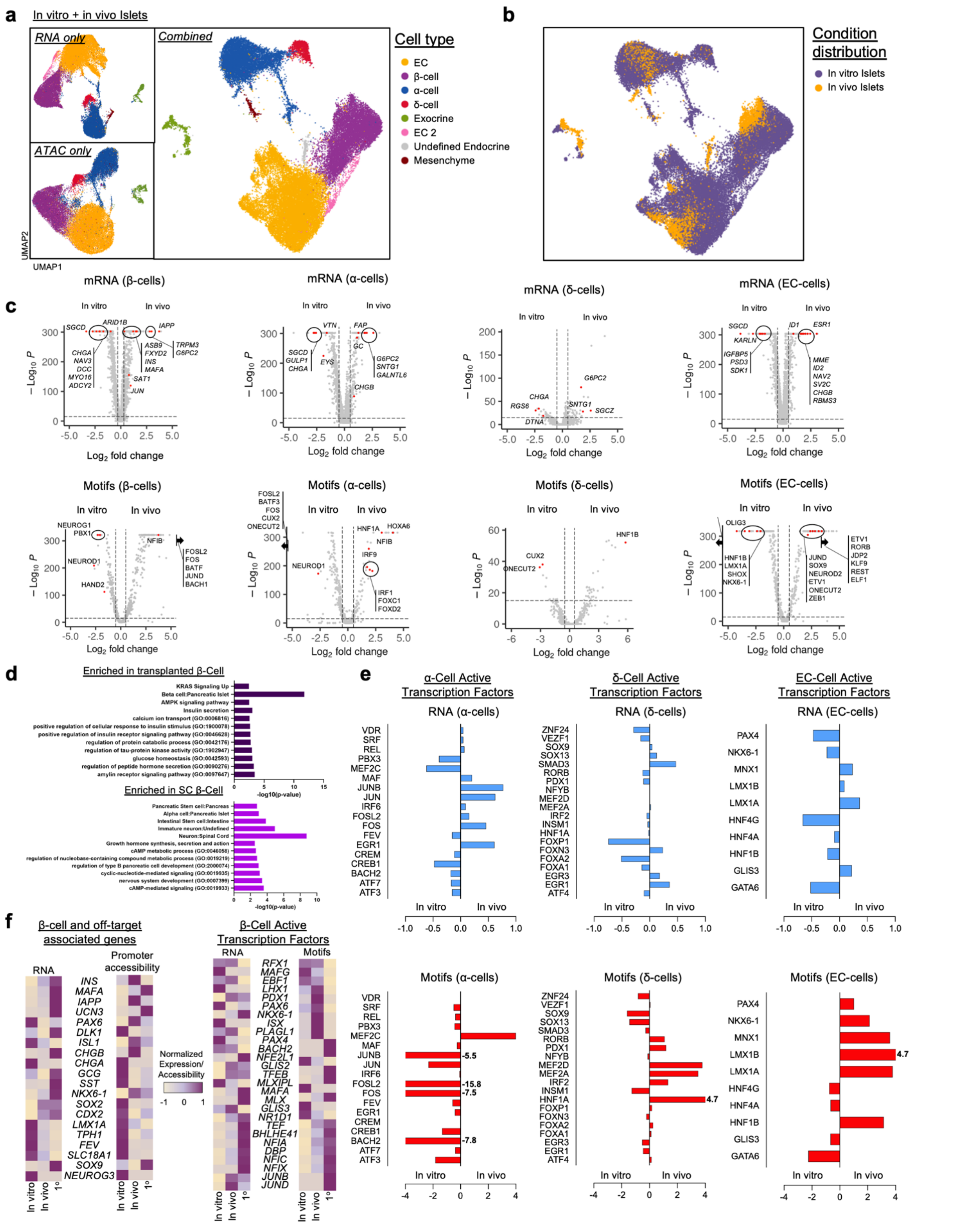


**Extended Data Fig. 8. | Multiomic sequencing comparison of in vitro SC-islets and transplanted in vivo SC-islets.** a, Integrative UMAP clustering showing cells from in vitro SC-islets and in vivo SC-islets using gene expression and chromatin information. b, UMAP showing distribution of SC-islet cells from in vitro or after transplanted in vivo condition. This plot highlights the separation of clusters from transplanted in vivo SC-β and SC-EC cells. c, Differential gene expression analysis (top) and motif chromatin accessibility analysis (bottom) of SC-β, SC-α, SC-δ and SC-EC cell populations from in vitro SC-islets and in vivo SC-islets. d, Gene set enrichment analysis showing enrichment of gene sets comparing in vitro SC-β cells and vivo SC-β cells. e, Bar graphs showing fold change differences of cell type associated active transcription factors in SC-α, SC-δ and SC-EC cells from in vitro and in vivo conditions. f, Heatmap comparing gene expression, promoter accessibility, or motif chromatin accessibility in β-cells from in vitro SC-islets, in vivo islets, and primary islets.

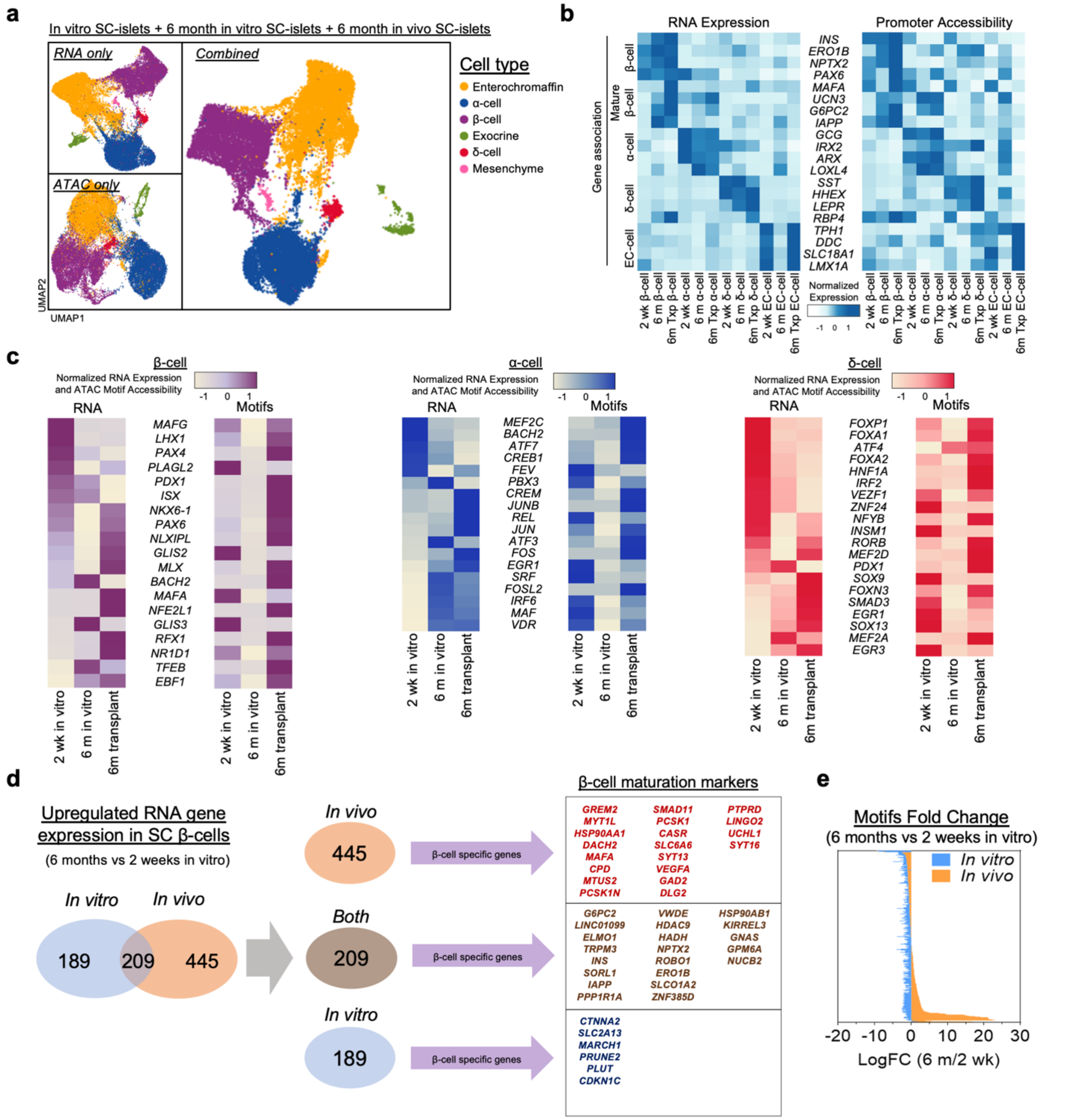


**Extended Data Fig. 9. | Multiomic sequencing comparison of week 2 in vitro SC-islets with 6 months in vitro SC-islets and 6 months in vivo SC-islets.** a, Integrative UMAP clustering showing cells from 2 weeks in vitro SC-islets, 6 months in vitro SC-islets, and 6 months in vivo SC-islets using gene expression and chromatin information. b, Heatmap showing gene expression and promoter accessibility of gene markers for SC-β, SC-α, SC-δ, and SC-EC cells. c,

Gene expression and motifs accessibility of active transcription factors associated with SC-β, SC-α, and SC-δ identity. d, Gene list showing markers for β cell identity 6 months in vitro, in vivo or both. Initial number of genes represent upregulated genes in the 6 month conditions (vitro and vivo) when compared to 2 week SC-islets. List of genes was cross-referenced with β cell upregulated genes from primary human islets. e, Plot highlighting greater increase of motifs accessibility from 6 months in vivo SC-β cells when compared in 6 months in vitro.

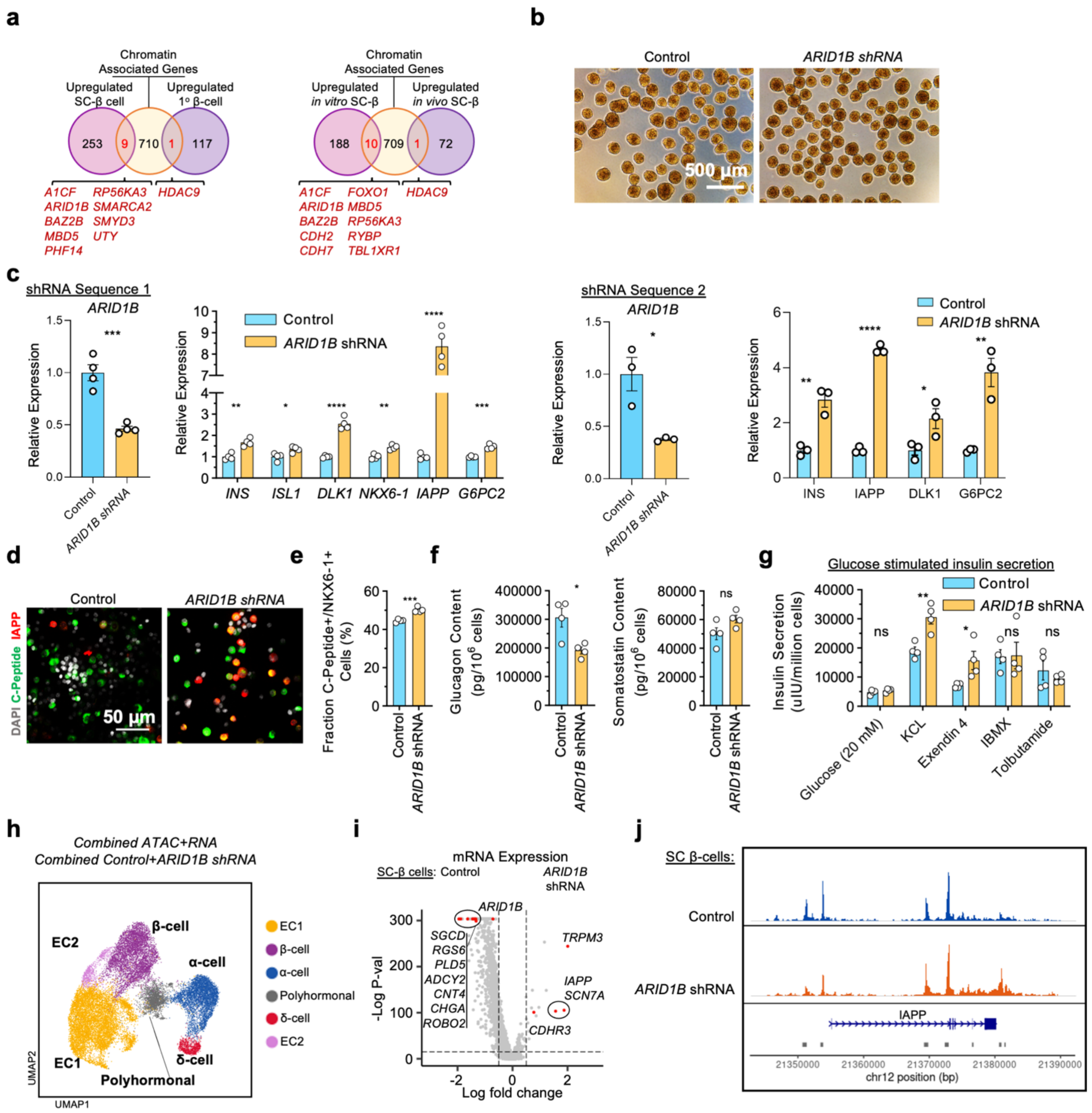


**Extended Data Fig. 10. | *ARID1B* knockdown increases expression of identity genes and chromatin features in SC-β cells.** a, Cross reference map of upregulated SC-β cell and primary β-cell genes (left) or transplanted SC-β cell genes (right) with chromatin associated genes to highlight regulators associated with each cell states. b, Brightfield images of SC-islets transfected with lentivirus carrying ARID1B shRNA. c, qPCR analysis, plotting mean ± s.e.m. (n

= 3-4), showing reduced expression of *ARID1B* using lentiviruses resulting in increased β cell identity gene expressions. (shRNA sequence 1: ARID1B, P = 5.3 x $10^{-4}$; *INS*, P = 0.0014; *ISL1*, P = 0.022; *DLK1*, P = 4.3 x $10^{-5}$; *NKX6-1*, P = 0.0015; *IAPP*, P = 3.0 x $10^{-6}$; *G6PC2*, P = 2.1 x $10^{-4}$) Two shRNA sequences were tested for validation of results. (shRNA sequence 2: ARID1B, P = 0.018; *INS*, P = 0.0035; *IAPP*, P = 5.9 x $10^{-6}$; *DLK1*, P = 0.045, G6PC2, P = 0.0054) d, Confocal fluorescent images showing increased expression of amylin in SC-islets with *ARID1B* knockdown. e, Flow cytometry analysis, plotting mean ± s.e.m. (n = 4), of SC-islets with *ARID1B* shRNA showing increased fraction of cells with C-peptide expression (P = 2.2 x $10^{-5}$) f, ELISA quantification, plotting mean ± s.e.m. (n = 4) of glucagon (P = 0.017), and somatostatin (ns, P = 0.087) content. g, Glucose stimulated insulin secretion assay, plotting mean ± s.e.m. (by ELISA, n = 4), comparing insulin secretion at high glucose (20 mM) stimulation from control (GFP shRNA) and ARID1B shRNA SC-islets in presence of various secretagogues. (Glucose (20mM), P = 0.27; KCL, P = 0.049; Exendin 4, P = 0.035; IBMX, P = 0.94; Tolbutamide, P = 0.43) h, Single-cell multiomic sequencing UMAPs of *ARID1B* knockdown SC-Islets, showing identified cell types, and cell knockdown condition using both gene expression and ATAC information. i, Differential gene expression analysis of SC-β cells comparing control and *ARID1B* shRNA. j, Chromatin accessibility around the *IAPP* genomic region of SC-β cells, showing increased peak signals with *ARID1B* shRNA. EC, enterochromaffin; ns, not significant.

### Supplementary Tables

#### Supplementary Table 1. Materials and methods

This table lists the reagents, cell lines, media formulations, and sequences used in this study. (1.1) Human stem cell lines used. (1.2) Details of the differentiation protocol to generate SC-islets. (1.3) Base medium formulations. (1.4) Factors and compounds used. (1.5) guide RNA sequences. (1.6) shRNA sequences. (1.7) Real-time PCR primer sequences.

#### Supplementary Table 2. Multiome sequencing details

This set of tables provides multiomic sequencing details such as individual data set details, cell filtering parameters, and total analyzed cell numbers. (2.1) Data set information used in this study. (2.2) Donor information details for adult human primary islet data sets. (2.3) Cell filtering paraments to exclude low quality cells, doublets, dead cells, and mouse cells from every data set. (2.4) Number of filtered cells from each data set used for analysis in this study.

#### Supplementary Table 3. Differential gene expression analysis for cluster identification.

This set of tables show gene expression analysis for cluster identification for SC-islets, primary islets, transplanted SC-islets, and CTCF CRISPRa SC-islets. (3.1) SC-islets. (3.2) Primary human islets. (3.3) Transplanted SC-islets. (3.4) CTCF CRISPRa SC-islets.

#### Supplementary Table 4. Differential motifs chromatin accessibility analysis for cluster identification.

This set of tables show motifs chromatin accessibility analysis for cluster identification for SC-islets, primary islets, transplanted SC-islets, and CTCF CRISPRa SC-islets. (4.1) SC-islets. (4.2)

Primary human islets. (4.3) Transplanted SC-islets. (4.4) CTCF CRISPRa SC-islets. Foldchange of motifs chromatin accessibility and RNA expression of cell type over other endocrine populations to determine transcriptional activity. (4.5) SC-islets TF Activity. (4.6) Primary islets TF Activity. (4.7) Transplanted SC-islets TF Activity.

**Supplementary Table 5. Trajectory analysis for delineation of SC-β cell and SC-EC cell population.**

This set of tables provides the source data for the multiomic sequencing analysis comparing SC-β cell and SC-EC cell populations. (5.1) Genes (mRNA) and values from trajectory analysis using Monocle. (5.2) Motifs accessibility and values from trajectory analysis using Monocle. (5.3) Differential gene (mRNA) expression analysis comparing SC-β cell and SC-EC cell. (5.4) Differential motifs chromatin accessibility analysis comparing SC-β cell and SC-EC cell.

**Supplementary Table 6. Cluster identification and characterization of subpopulations.**

This set of tables show differential gene expression and motifs accessibility analysis of subpopulations in SC-β cells, SC-EC cells, and primary β cells. (6.1) SC-β subpopulation 1 (6.2) SC-β subpopulation 2 (6.3) SC-β subpopulation 3 (6.4) SC-β subpopulation 4 (6.5) SC-EC subpopulation 1 (6.6) SC-EC subpopulation 2 (6.7) SC-EC subpopulation 3 (6.8) SC-EC subpopulation 4 (6.9) Primary β subpopulation 1 (6.10) Primary β subpopulation 2 (6.11) Primary β subpopulation 3.

**Supplementary Table 7. Differential analysis of CTCF CRISPRa experiments.**

This set of tables provides the source data for the multiomic sequencing analysis of CTCF overexpression in SC-islets. (7.1) Differential gene expression and motifs chromatin accessibility analysis comparing SC-EC cells and SC-iEC cells. (7.2) Differential gene expression and motifs chromatin accessibility analysis comparing the SC-endocrine population from control and CTCF over expression.

**Supplementary Table 8. Differential gene expression, motifs accessibility and promoter accessibility analysis of SC-islets and primary islets.** This set of tables provides differential analysis comparing SC-islet cells with primary islet cells (8.1) SC-β cells vs primary β cells. (8.2) SC-α cells vs primary α cells. (8.3) SC-δ cells vs primary δ cells.

**Supplementary Table 9. Differential gene expression, and motifs accessibility analysis of short-term and long-term in vitro SC-β cells.** This set of tables provides differential analysis comparing short term in vitro (week 2 – 4) with long term (month 6 -12) SC-β cells (9.1) Differential gene expression. (9.2) Differential motifs accessibility.

**Supplementary Table 10. Differential gene expression, motifs accessibility and promoter accessibility analysis of SC-islets and primary islets.** This set of tables provides differential analysis comparing SC-islet cells with transplanted SC-islet cells. (10.1) SC-β cells vs transplanted SC-β cells. (10.2) SC-α cells vs transplanted SC-α cells. (10.3) SC-δ cells vs transplanted SC-δ cells. (10.4) SC-EC cells vs transplanted SC-EC cells.

**Supplementary Table 11. Differential gene expression and motifs accessibility analysis comparing 6 months in vitro SC-islets with 6 months in vivo SC-β cells.** This set of tables provides differential analysis comparing 6 months in vitro SC-β cells with transplanted SC-β cells. (11.1) 6 months in vitro SC-β cells vs 6 months in vivo (transplanted) SC-β cells.

**Supplementary Table 12. Differential gene expression and motifs accessibility analysis comparing SC-β cells from control and ARID1B shRNA condition.** This set of tables provides differential analyses comparing control SC-β cells with ARID1B shRNA SC-β cells. (12.1) Control (GFP shRNA) SC-β cells vs ARID1B shRNA SC-β cells.
