## Supplementary Figure for "Defining the chromatin and transcriptional landscape of stem cell-derived islets"

Supplementary Figure 1

#### Cell exclusion strategy for single-cell multiome analysis

##### SC-Islet and human islet

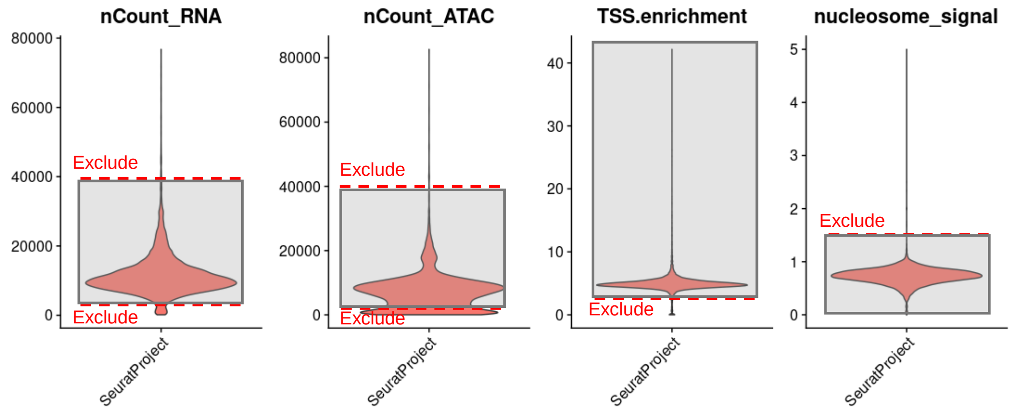


##### Transplanted SC-islet

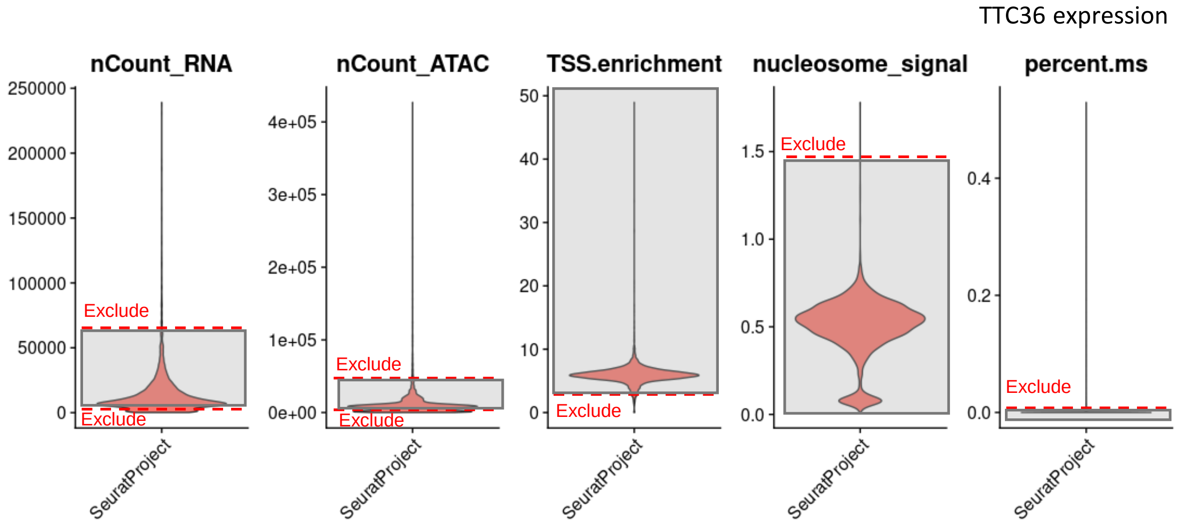

Supplementary Figure 2

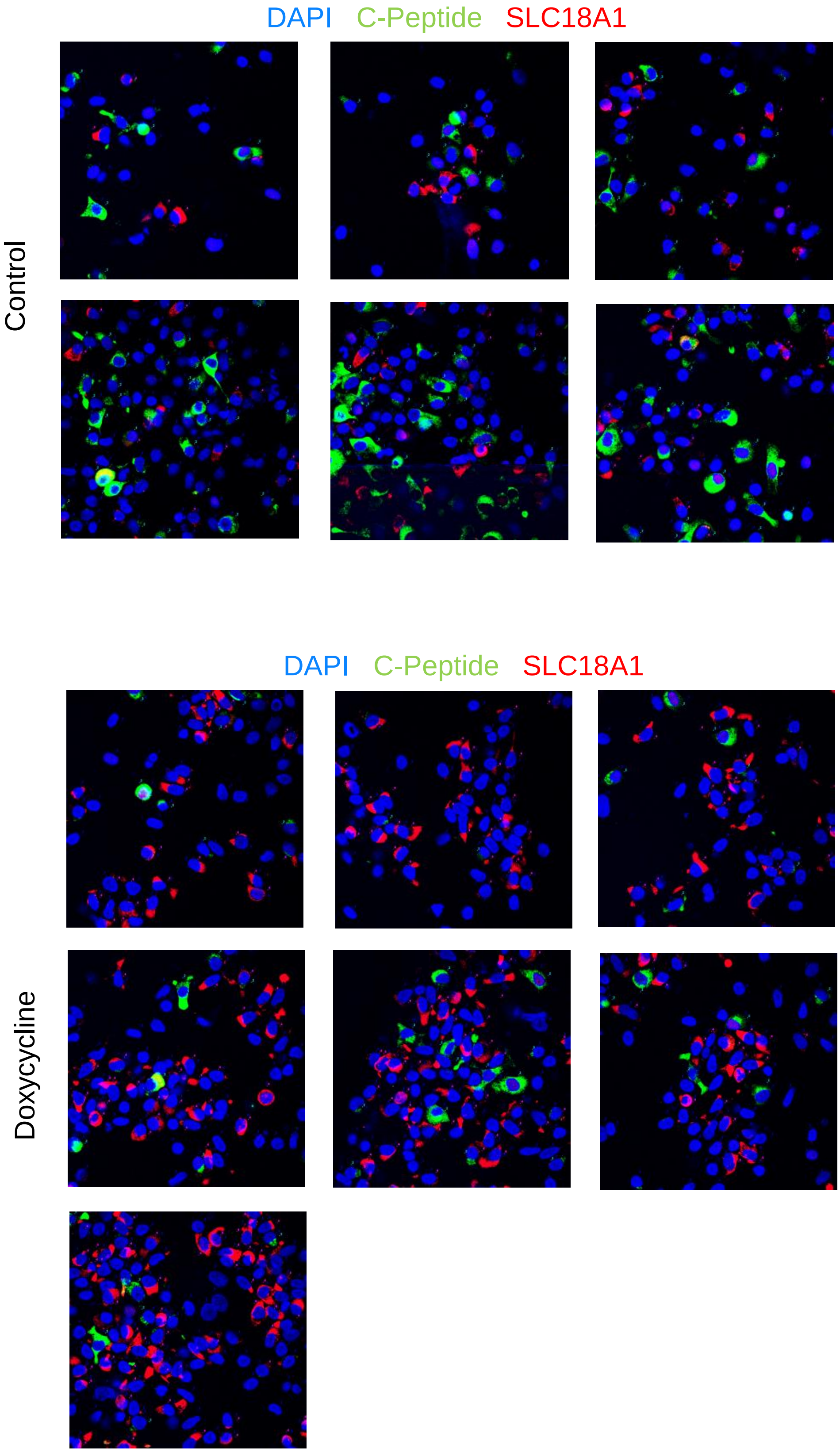

Supplementary Figure 3

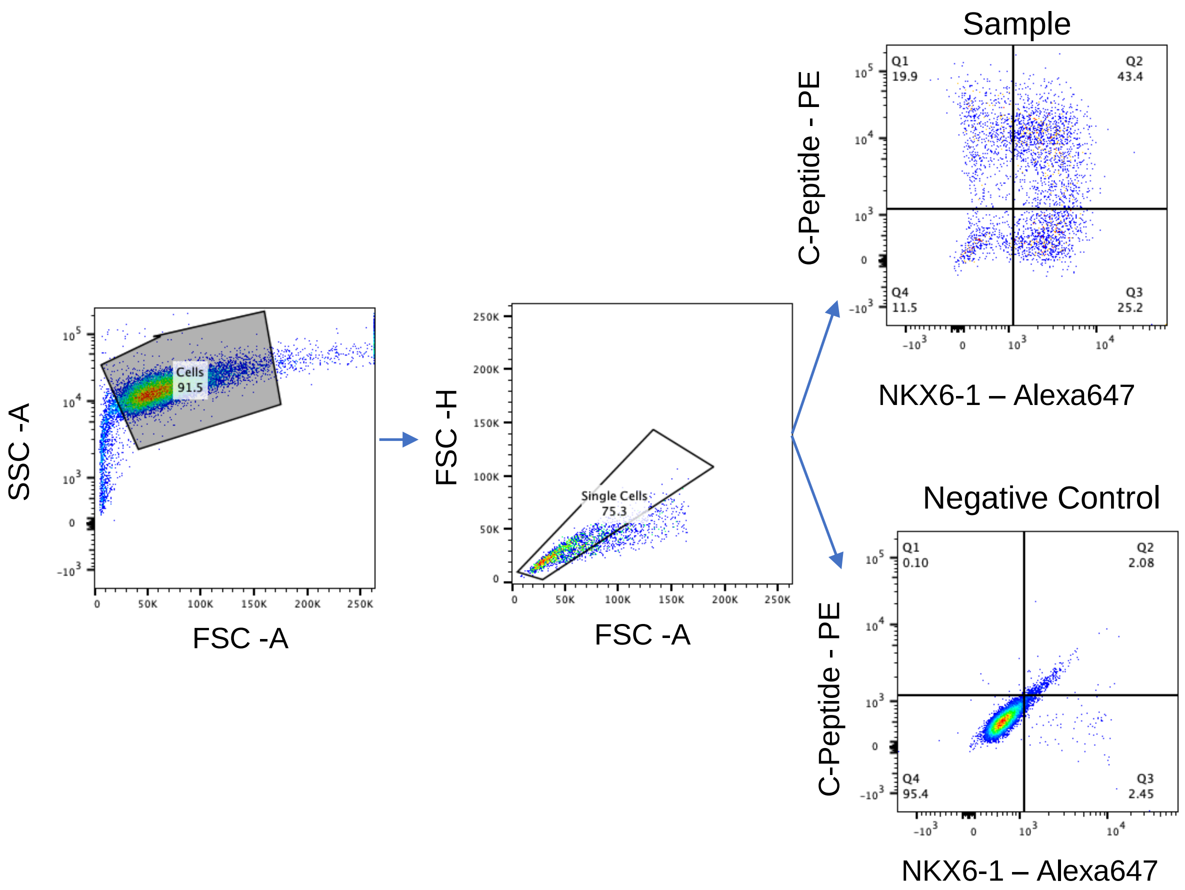

### Supplementary Figure 4

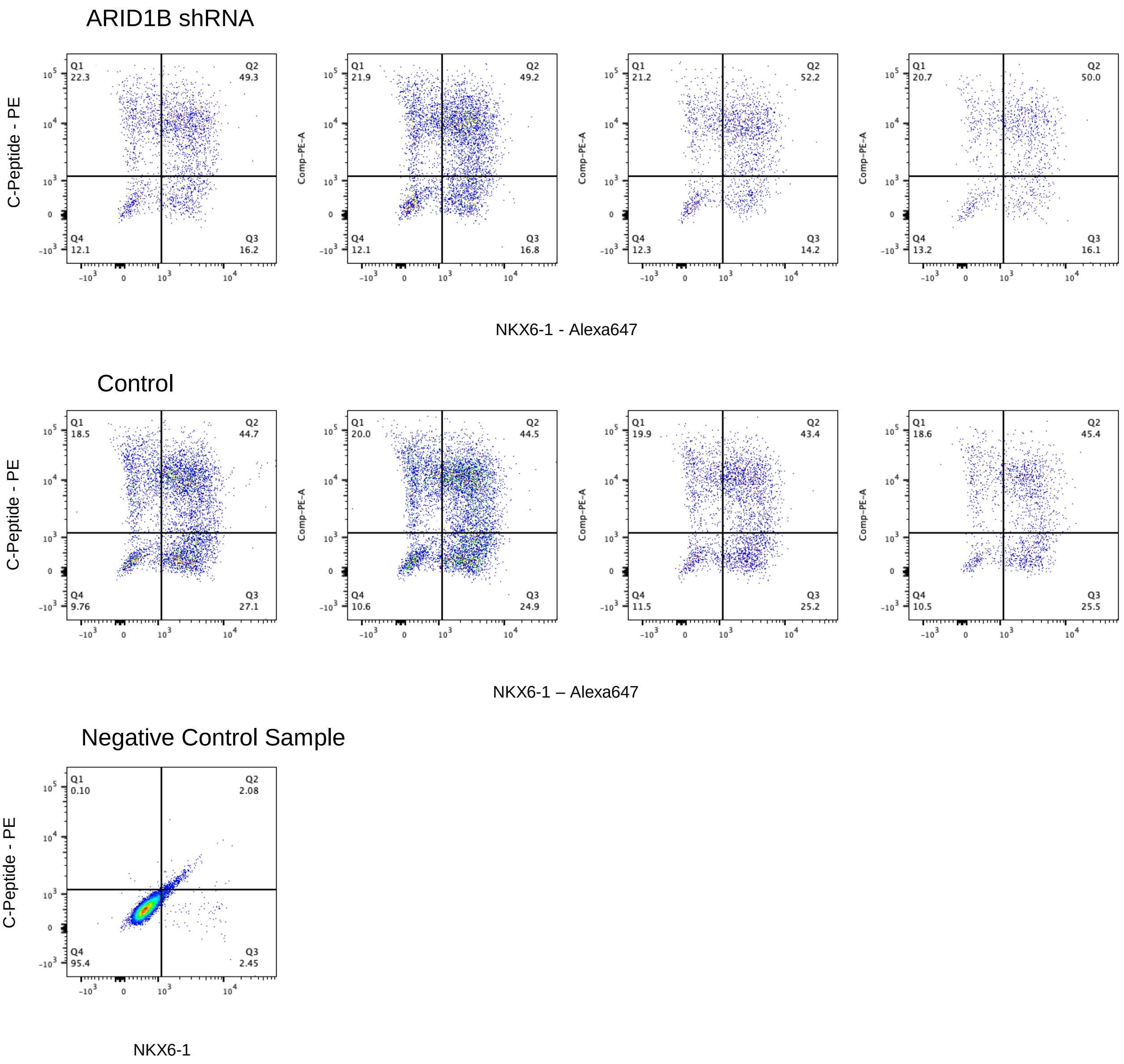
